# Composition of the North American wood frog (*Rana sylvatica*) skin microbiome and seasonal variation in community structure

**DOI:** 10.1101/2020.01.28.921544

**Authors:** Alexander J. Douglas, Laura. A. Hug, Barbara A. Katzenback

**Affiliations:** Department of Biology, University of Waterloo, Waterloo, Ontario, Canada, N2L 3G1

**Keywords:** Microbiome, Amphibian, *Rana sylvatica*, Skin, Innate Immunity, Season

## Abstract

While a number of amphibian microbiomes have been characterized, it is unclear how microbial communities might vary in response to seasonal changes in the environment and the behaviors which many amphibians exhibit. Given recent studies demonstrating the importance of the skin microbiome in frog innate immune defenses against pathogens, investigating how changes in the environment impact the microbial species present, and thus their potential contribution to skin host defense, will provide a better understanding of conditions that may alter host susceptibility to pathogens in their environment. We sampled the skin microbiome of North American wood frogs (*Rana sylvatica*) from two breeding ponds in the spring, along with the microbial community present in their vernal breeding pools, and frogs from the nearby forest floor in the summer and fall to determine whether the microbial composition differs by sex, vernal pond site, or temporally across season (spring, summer, fall). Taxon abundance data reveals a profile of bacterial phyla similar to those previously described on anuran skin, with Proteobacteria, Bacteroidetes, and Actinobacteria dominating the wood frog skin microbiome. Our results indicate that sex had no significant effect on skin microbiota diversity, however, this may be due to our limited female sample size. Vernal pool site had a small but significant effect on skin microbiota, but skin-associated communities were more similar to each other than to the communities observed in the frogs’ respective pond water. Across seasons, diversity analyses suggest there are significant differences between the skin microbiome of frogs from spring and summer/fall groups while the average α-diversity per frog remained consistent. Bacterial genera known to have antifungal properties such as *Pseudomonas* spp. and *Rhizobium* spp. were prevalent, and several were considered core microbiota during at least one season. These results illustrate seasonal variation in wood frog skin microbiome structure and highlight the importance of considering temporal trends in an amphibian microbiome, particularly for species whose life history requires recurrent shifts in habitat and behavior.

## 1 Introduction

Amphibians have unique communities of skin-dwelling microbes regulated by various mechanisms, including inoculation from the environment, skin-sloughing and specific skin secretions [1, 2]. Studies of the amphibian skin microbiome have revealed that while many bacterial phyla are present, species from Acidobacteria, Actinobacteria, Bacteroidetes, Cyanobacteria, Firmicutes and Proteobacteria tend to dominate [1, 3–5]. Many families and genera within these phyla are present on a wide range of amphibian species, but the specific species tend to be unique to the host. Additionally, species which dominate the skin-associated community are not generally those which are dominant in the environment at large, and therefore must be selected for by the skin microenvironment [1]. Further microbiome studies have shown that frogs of the same species sampled during different seasons or from different habitats can have large differences in the diversity of bacteria present on their skin [4, 6, 7], underscoring the influence of environmental factors on the microbial community. Most studies of amphibian microbiota have been performed on amphibian species from tropical or subtropical climates, meaning data on temperate amphibian species is less comprehensive. Therefore, it is currently unclear how stronger seasonal differences in climate and host behavior might influence the amphibian skin microbiome.

Although the functional significance of amphibian skin microbiota are not yet well defined, recent studies have highlighted the importance of the skin commensal microbiome to the amphibian host skin defense mechanisms against invading pathogens [2, 8–10]. Through the production of antifungal metabolites [11], some bacterial species commonly found on the skin of amphibians inhibit the growth of the pathogenic fungi *Batrachochytrium dendrobatidis* [12, 13], the causative agent of amphibian chytridiomycosis and proximate cause of amphibian declines on multiple continents [14, 15]. Transfer of these specific bacteria to the skin of salamanders suffering from chytridiomycosis reduced the severity of symptoms [16], which made a strong case for the protective effects of select commensal bacterial species and has spurred the use of bacterial species as probiotic washes for target amphibian populations [17]. However, the effectiveness of specific bacteria to inhibit infection has been shown to vary based on factors such as the strains of the symbiont and pathogen as well as temperature [18], so it is unlikely that any single species of bacteria would provide universal protection. Additionally, pathogen infection drives changes in the structure of bacterial communities on the skin and correlates with lower community diversity [19–21]. It is increasingly clear that the characteristics of the amphibian skin microbiome are related to infection status and that the structure and diversity of the commensal microbial community may be key to predicting pathogen resistance. Thus, understanding how frog skin microbial community composition varies across environmental conditions would provide important information in predicting environments where amphibians may be more or less susceptible to pathogens.

The North American wood frog (*Rana sylvatica*) is widespread with a range extending through most of Canada, Alaska, and the Northeastern United States [22]. *R. sylvatica* inhabits uplands environments and the far north where few, if any, other frog species inhabit. Wood frogs breed in temporary pools in the early spring, leave these pools to migrate upland into the terrestrial forest environment during warmer months [23] and hibernate on the forest floor in winter, routinely surviving multiple sustained bouts of whole body freezing [24, 25]. *R. sylvatica* is susceptible to infection by Frog Virus 3 and *B. dendrobatidis* [26, 27], pathogens which threaten amphibian populations worldwide. Despite the wide range of *R. sylvatica* and its known susceptibility to pathogens of concern, no comprehensive study of skin microbiota has ever been performed on this species. Given the seasonal shifts in habitat and behavior experienced by *R. sylvatica*, they are well-suited to investigate trends which may apply to other temperate species, such as whether seasonal changes in microbiome composition might lead to windows of particular protection from, or susceptibility to, amphibian pathogens, as has been explored in other species [20, 28, 29].

The objectives of this study are to (1) identify the composition of the *R. sylvatica* skin microbiome and (2) determine whether the skin microbiome community structure varies with sex, vernal pool of origin or across seasons. It is hypothesized that the wood frogs carry a range of bacterial phyla similar to other frog skin microbiomes, with variation in representation and abundance of bacterial taxa between individuals to reflect separate ponds of origin and season of capture. To determine whether this is the case, we have analyzed the microbial community present on the skin of *R. sylvatica* using 16S rRNA gene amplicon sequencing. Skin swabs were obtained from frogs captured from two spatially separated ponds in the spring, and the surrounding forest in summer and fall so that skin-associated microbial communities might be compared on a seasonal basis, as well as between different capture sites.

## 2 Materials and Methods

### 2.1 Experimental Design & Sample Collection

Wild *R. sylvatica* adults were sampled from a site in the Waterloo Region of Ontario, Canada, during the spring (April - May), summer (July - August) and fall (October). In the spring, individuals were captured from two vernal ponds spatially separated by ∼200 m, herein referred to as Pond 1 and Pond 2, while individuals were captured from the surrounding forest floor during the summer and fall seasons. Frog skin microbiota sample sizes were dependent on the number of frogs that could be captured successfully. Sampling numbers are given for each site and season in Table 1.

**Table 1.** Summary of frog skin swab samples. Season, site and sex of frog are given for each skin swab sample. Male frogs are denoted with an “M”, while females are denoted with an “F”

| Season | Capture Site |  |  |
| --- | --- | --- | --- |
|  | Pond 1 | Pond 2 | Forest Floor |
| Spring 2018 | 13 M / 2 F | 8 M / 4 F |  |
| Summer 2018 |  |  | 11 M / 0 F |
| Fall 2018 |  |  | 5 M / 2 F |
| Spring 2019 | 10 M / 0 F | 12 M / 0 F |  |

Individuals were captured by nets and each frog was handled with a new pair of sterile nitrile gloves. Wood frogs were gently rinsed with sterile distilled water to remove transient microbes. To collect resident microbes frogs were swabbed with a sterile rayon-tipped applicator (Puritan Medical Products Company, LLC., Guilford, ME, USA) 12 times on both the dorsal and ventral surfaces, covering as much of the skin surface as possible. To control for environmental microbes that may have deposited onto the swab during the sampling process, “field control” swabs were produced by wetting a clean rayon swab with sterile distilled water and mimicking the swabbing action in the open air. Each rayon swab head was placed into a sterile 1.7 mL microfuge tube and the applicator stick cut just above the rayon tip using flame-sterilized scissors. Samples were transported on ice prior to storage at −80 °C. Animal care and handling was performed in accordance with the guidelines of the University of Waterloo Animal Care Committee and the Canadian Council on Animal Care (Animal Utilization Projects #30008 and #40721), and animals captured under the Ontario Ministry of Natural Resources and Forestry Wildlife Scientific Collectors Authorization Permits (#1088586 and 1092603) issued to Dr. B.A. Katzenback.

When individuals were captured from vernal pools, a water sample was taken by pushing 50 mL of pond water, taken from just below the surface of the water, through a Sterivex-GP PES 0.22 µm filter (Millipore, Burlington, MA, USA) using a 50 mL syringe (Fisher). The filter units were disconnected from the syringe, placed in individual sterile 50 mL conical tubes (FroggaBio) and held on ice prior to storage at −80 °C.

### 2.2 DNA Isolation and Amplicon Sequencing

DNA isolation was performed using the DNeasy PowerSoil Kit (QIAGEN Inc., Venlo, Netherlands) according to the manufacturer’s protocol. Frog skin swab samples or field controls were removed from −80 °C and immediately transferred from their storage tubes into PowerBead tubes. DNA was isolated from vernal pool microbiota filtride by removing the filter paper from the cartridge and cutting the filter paper into thin strips using flame-sterilized scissors before addition to the PowerBead tubes. To control for bacterial contamination from the laboratory environment and/or the extraction kit components, a clean rayon swab was wet with sterile distilled water and cut into a labelled 1.7 mL microfuge tube with flame sterilized scissors to act as a “process control” and was processed alongside the samples. All samples (skin swabs, vernal pool filters, field controls, process controls) were immediately vortexed for 10 s after transfer to the PowerBead tubes prior to following the manufacturer’s protocol. Isolated DNA was eluted in the provided elution buffer and stored at −80 °C.

Presence of bacterial DNA was confirmed using PCR amplification. Each reaction contained 18.875 µL of molecular biology grade water, 2.5 µL of 10× PCR buffer, 0.5 µL of 10 µM dNTPs, 1 µL of 10 µM forward primer 515FB 5’-GTGYCAGCMGCCGCGGTAA-3’ [30, 31], 1 µL of 10 µM reverse primer 926R 5’-CCGYCAATTYMTTTRAGTTT-3’ [30, 32], and 0.125 µL Taq DNA polymerase (5 units/µL) (GeneDirex) per 1 µL of sample, for a total reaction volume of 25 µL. PCR was performed with the following cycling conditions: 94 °C for 3 min, then 35 cycles of 94 °C for 45 s, 50 °C for 1 min and 72 °C for 1.5 min, followed by extension at 72 °C for 5 min. PCR product were separated by electrophoresis at 130 V for 30 min on 2% agarose gel in TAE buffer and visualized using SafeRed dye and trans-UV (302 nm) imaging in a ChemiDoc XRS+ (Bio-Rad).

Samples were sent for 16S rRNA gene amplicon sequencing by MetagenomBio (Waterloo, ON, Canada). PCR reactions were prepared in triplicate for each sample. Each 25 µL reaction mixture contained 1.6 µL of molecular grade water, 0.2 µL of BSA (20 mg/mL), 2.5 µL of 10× standard Taq buffer, 0.5 µL of 10 mM dNTPs, 5.0 µL of 1 µM forward primer 515FB 5’-GTGYCAGCMGCCGCGGTAA-3’ [30, 31], 5.0 µL of 1 µM reverse primer 806RB 5’-GGACTACNVGGGTWTCTAAT-3’ [30, 31], 0.2 µL of Taq DNA polymerase (5 units/µL), and 10 µL of sample DNA. PCR was performed with the following thermocycling conditions: 95°C for 5 min, 35 cycles of 95°C for 30 s, 50°C for 30 s and 72°C for 60 s, followed by an extension at 72°C for 10 min. The products of the triplicate reactions were pooled and resolved with 2% TAE agarose gel. PCR products of the correct amplicon size (∼ 291 bp) were excised, pooled, gel purified and quantified using the Invitrogen™ Qubit™ dsDNA HS Assay Kit (Thermo Fisher Scientific Inc., Waltham, MA, USA). For all 2018 samples sequencing was performed using an Illumina MiSeq and the MiSeq Reagent Kit v2 (Illumina, Inc., San Diego, CA, USA) for 2 sets of 250 cycles. This was increased to 3 sets of 250 cycles for all 2019 samples due to an error at the sequencing center

### 2.3 Amplicon Sequence Data Processing

Sequence data was obtained as FASTQ files in the CASAVA 1.8 paired-end demultiplexed format. Files from repeat sequencing runs were concatenated to create a FASTQ file containing all of the observed sequences for each sample. These files were imported into QIIME 2 v2019.1.0 [33] and all analysis was performed using QIIME 2 unless otherwise stated.

Using DADA2 [34], reads were trimmed by 25 bp on the 5’ end to remove the primer sequence and truncated to 245 bp to remove low quality regions, filtering out any reads shorter than this length. The reads were dereplicated, denoised and any chimeric sequences were removed. Paired forward and reverse reads were merged, generating the final amplicon sequence variants (ASVs). Each ASV was assigned taxonomy using a naïve Bayesian classifier trained on the SSU Ref NR 99 dataset from the SILVA 132 release [35], with sequences trimmed to include only the V4-V5 region. All unassigned ASVs and those assigned to chloroplast or mitochondria were filtered from the samples. To minimize erroneous ASVs, a minimum frequency of 22 (0.001% of the total sequence count) was set, and any ASVs with a total frequency less than 22 were filtered from the samples.

### 2.4 Taxonomic Assignment and Significance

A multiple sequence alignment (MSA) was produced from the ASVs using MAFFT [36]. Columns of the alignment which were ambiguously aligned were masked to avoid introducing error to the phylogenetic model. A phylogenetic tree was generated from the MSA using RAxML [37] with 100 bootstraps. The RAxML tree was assigned a midpoint root and used for all further phylogenetic analysis.

Core taxa were determined for each group of seasonal frog skin communities, and for the overall frog skin microbiome, using the core-features command in QIIME2. Core taxa were defined as those found to be present in 90% or more of the samples within a given group, as this is a commonly used threshold [6, 38] which meets the definition of core microbiota as those which are commonly present within samples from a given environment [39]. Venn diagrams depicting the overlap of seasonally core microbiota were prepared using the online Venn diagram tool provided by the University of Ghent (http://bioinformatics.psb.ugent.be/webtools/Venn/).

To assess the presence of microbiota with putative antifungal activity, bacterial taxa identified from the *R. sylvatica* skin microbiome were searched against the Antifungal Isolates Database [40]. The metadata file for the Antifungal Isolates Database was obtained and sorted based on genera, filtering for those genera which were detected on *R. sylvatica* skin. Genera with one or more antifungal isolates were considered putatively antifungal, and their relative abundances in the wood frog skin microbial community were compared between seasons.

### 2.5 Statistical Analyses

For diversity analyses, samples were rarified to 10,000 sequences. Those containing less than 10,000 were omitted from diversity analyses. The phylogenetic tree and rarified ASV frequencies were used to calculate various α-diversity metrics (Faith’s Phylogenetic Diversity, Shannon’s Diversity Index, ASV richness). Diversity was compared between groups using a one-way analysis of variance (ANOVA).

To assess differences in microbial community composition between samples, various β-diversity metrics were calculated (Unweighted UniFrac, Weighted UniFrac and Bray-Curtis dissimilarity). β-diversity was visualized using principal coordinate analysis (PCoA). The *adonis* function from the R vegan package was used to perform permutational multivariate analysis of variance (PERMANOVA) with 999 permutations. To determine whether groups had significant differences in microbiome composition, pairwise PERMANOVA tests were applied to the dissimilarity matrix produced by each β-diversity metric and performed with 999 permutations.

## 3 Results

### 3.1 Microbiome Overview Statistics

We obtained microbial 16S rRNA amplicon sequences from each of the 66 frog swabs, 5 water samples, 5 field blanks, and 3 process blanks. After filtering based on quality, taxonomy and minimum frequency, a total of 1,937,866 sequences remained. Sequence counts varied between samples considerably, and those samples sequenced in the later set (Spring 2019 frog swabs, water samples and associated blanks) had much higher counts on average due to their increased sequencing depth. A total of 4,325 ASVs were recognized, which ranged in frequency from 22 to 159,313 appearances, with a median frequency of 60. These ASVs were matched to 1,384 unique taxonomic assignments, 1,123 of which were specific to at least the genus level.

We found that DNA isolated from process controls yielded ASVs belonging to 9 bacterial phyla and contained 37 ASVs each on average, which equated to a total of 71 unique taxa when combined at the genus level (**Supplementary Table 1**). Unlike the communities found on frog skin and in pond water, the majority of ASVs found in the process controls belonged to a limited group of genera, primarily *Curtobacterium* (43%), with *Lactobacillus* (16%) and *Acholeplasma* (7.7%) also more abundant. The field controls generally had more diverse communities, with an average of 64 ASVs per blank. These ASVs belonged to a total of 15 phyla, and 164 unique taxa when combined at the genus level (**Supplementary Table 2**). The genera with the greatest mean relative abundance was once again *Curtobacterium* (8.4%), but genera which were abundant on frog skin such as *Massilia* and *Pseudomonas* were also found to be abundant in the field blanks (4.1% and 3.8%, respectively). On average 11% of amplicons detected in frog skin samples and 0.07% of amplicons detected in water samples were ASVs which were also found in the process controls, while an average of 35% of amplicons detected in frog skin samples and 32% of amplicons detected in water samples were ASVs which were also found in the field controls. Of the 11% of the amplicons found on frogs which were shared with the process blank, the majority (5.8% of mean relative abundance) corresponded to a single ASV belonging to *Curtobacterium*. A large portion of the ASVs belonging to *Curtobacterium* were found on a group of five frogs where this ASV made up more than 50% of amplicons detected.

### 3.2 Capture Site Influences the Structure of Microbial Communities

Spring was the only season in which sampling occurred across two distinct sites, the two vernal pools. It was thus necessary to investigate how pond of origin might influence an individual’s skin microbiome. We observed that the skin microbiota of wood frogs captured from Pond 1 and Pond 2 had very similar α-diversity metrics, and both of the skin-associated communities had higher average microbial diversity than the microbial communities present in the water samples taken from either vernal pond (Fig. 1). When we sorted frog skin microbiota and vernal pond microbiota samples from spring 2018 and spring 2019 into groups based on their pond of origin, there were significant differences in α-diversity (ANOVA: ASV richness: p = 0.0017; Shannon diversity index: p = 0.0032; Faith’s phylogenetic diversity: p = 0.0041). Differences were between the Pond 2 water and the Pond 1 frog skin (pairwise ANOVA: ASV richness: p = 0.0355; Shannon diversity index: p = 0.0629; Faith’s phylogenetic diversity: p = 0.0068) and between Pond 2 water and Pond 2 frog skin (pairwise ANOVA: ASV richness: p = 0.0029; Shannon diversity index: p = 0.0114; Faith’s phylogenetic diversity: p = 0.0035). There were no significant differences in community diversity between Pond 1 water and Pond 1 frog skin, Pond 1 water and Pond 2 frog skin, Pond 1 water and Pond 2 water or Pond 1 frog skin and Pond 2 frog skin.

**Fig. 1.**
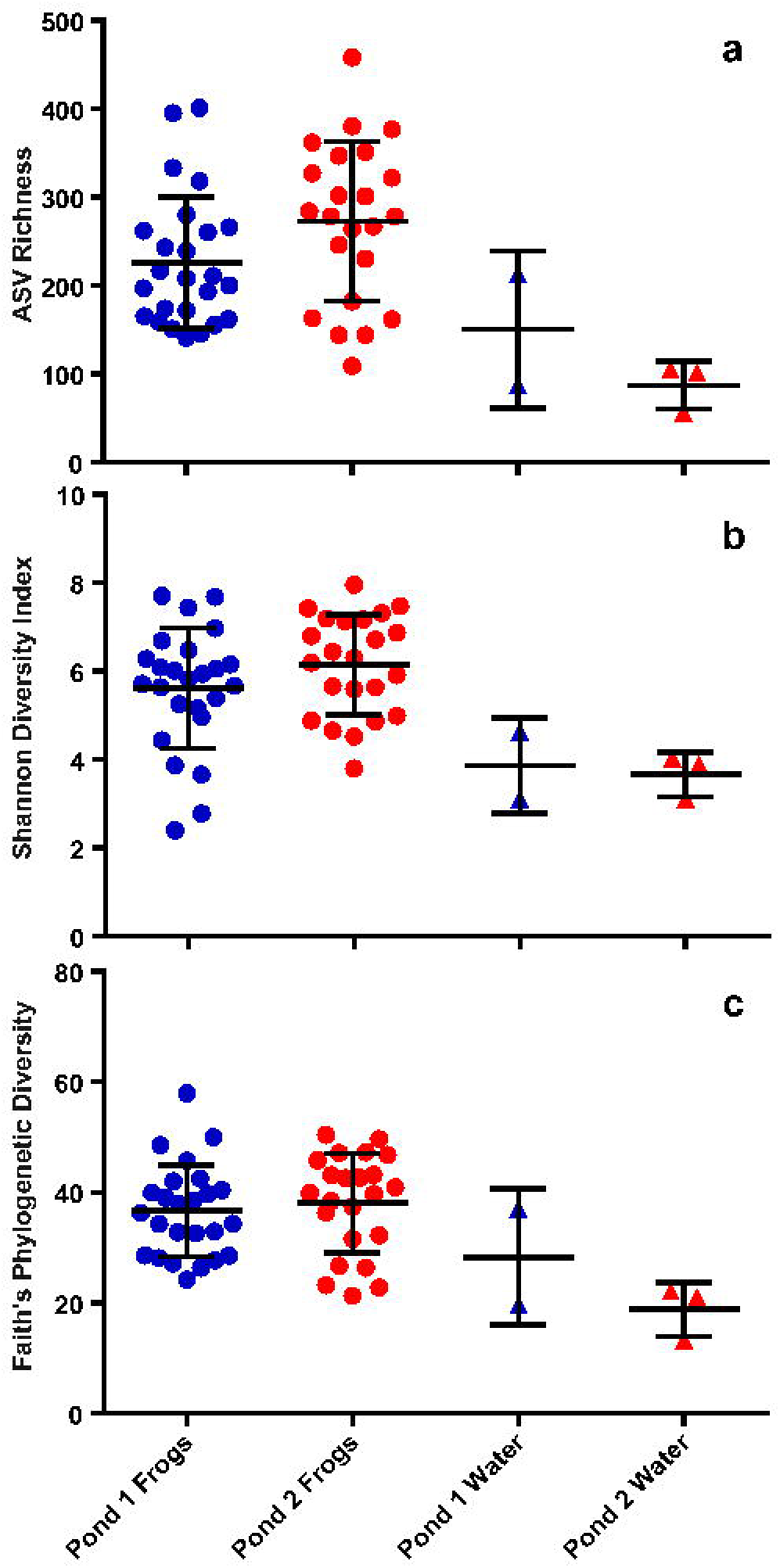
Comparison of frog swab and pond water α-diversity diversity metrics. Sample α-diversity was calculated using a sampling depth of 10,000. Mean seasonal value and standard deviation of each group is shown. Results are given for (**a)** ASV Richness, (**b)** Shannon’s Diversity Index and (**c)** Faith’s Phylogenetic Diversity

The microbial communities present in each group of frog skin samples and water samples included many of the same bacterial phyla and had a high proportion of ASVs belonging to Proteobacteria, Bacteroidetes and Actinobacteria (Fig. 2). These three phyla were the only phyla we found to be highly abundant in the water samples, while all other phyla present had a mean relative abundance of less than 1%. The frog skin communities from both ponds also had Acidobacteria and Verrucomicrobia ASVs present at greater than 1% mean relative abundance, and Pond 2 frog skin community additionally had Firmicutes ASVs present at more than 1% mean relative abundance. We employed pairwise analysis of variance (ANOVA) to compare the relative abundance of each of these 6 common phyla between the water samples and frog skin samples from both ponds and no significant difference in abundance was observed for any of the phyla. Beyond the most abundant phyla, the frog skin microbiota exhibited a more diverse range of phyla compared to vernal pool microbiota; the skin-associated communities from frogs captured from Pond 1 and Pond 2 included ASVs from phyla that were not present in any pond water samples.

**Fig. 2.**
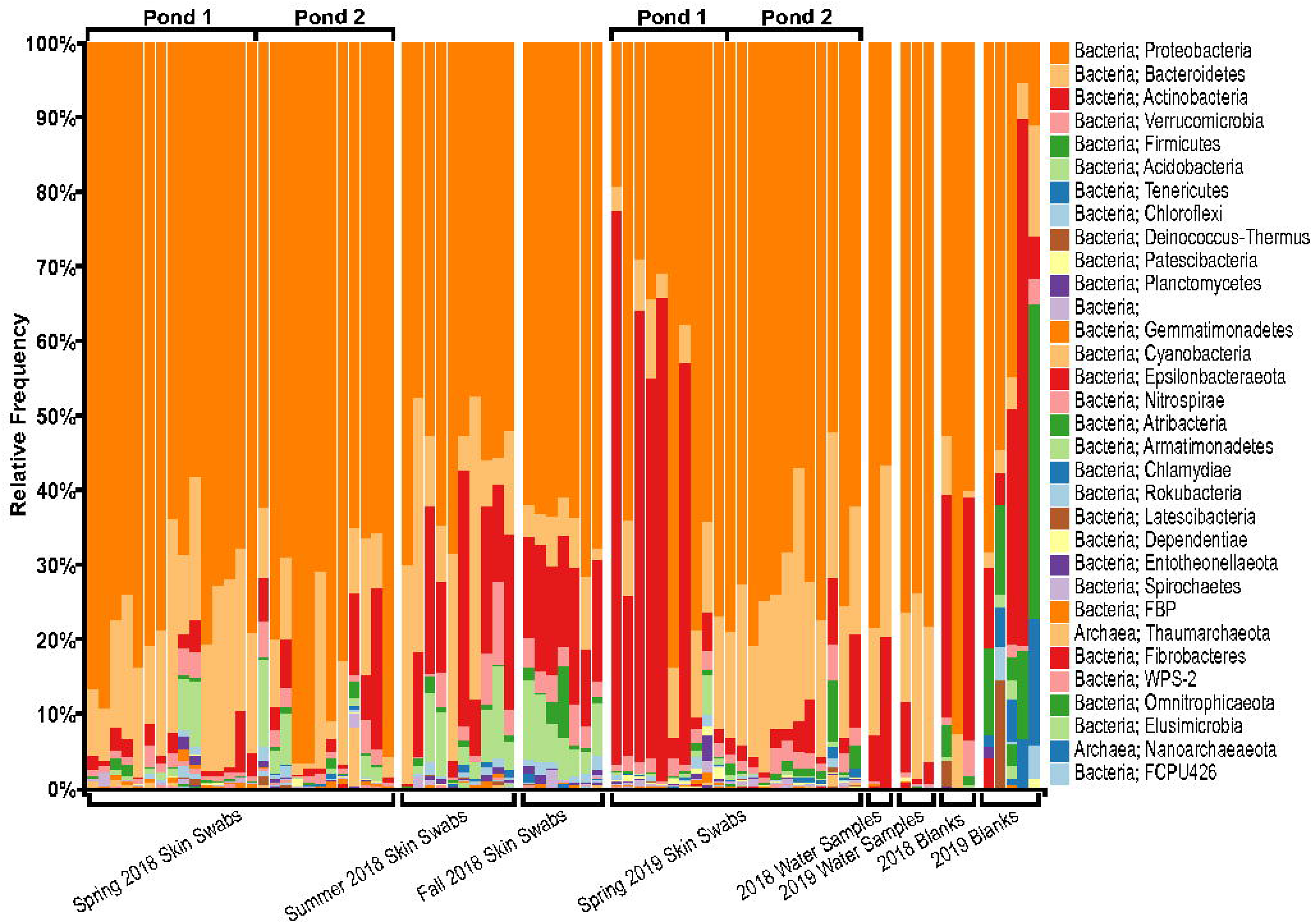
Relative frequency of microbial phyla ASVs present in individual *R. sylvatica* skin swab samples and controls. Phyla are listed from top to bottom in order of decreasing summed total ASV frequency. Bars represent relative frequency within a sample and are given in corresponding order

Sample type (frog skin microbiota, vernal pond microbiota), site (Pond 1, Pond 2) and year all had significant (p < 0.005) but small (R^2^ < 0.12) effects on community composition and abundance of the observed taxa when assessed using *adonis* (Table 2). Sample type had the largest effect, particularly when considering abundance as observed with Weighted UniFrac distance (Table 2b, *adonis* pseudo-F = 6.86, p < 0.001, R^2^ = 0.11) and Bray-Curtis dissimilarity (Table 2c, *adonis* pseudo-F = 7.71, p < 0.001, R^2^ = 0.12). Effect of the sample type (Table 2a, *adonis* pseudo-F = 4.08, p < 0.001, R^2^ = 0.07) and year (Table 2a, *adonis* pseudo-F = 4.20, p < 0.001, R^2^ = 0.07) had equal effects as observed with Unweighted UniFrac distance. Principal coordinate analysis of the β-diversity metrics did not reveal clear clustering of frog skin microbiota by pond origin under any conditions (Fig. 3). Water samples were found to cluster separately from frog samples only when relying on Bray-Curtis dissimilarity values (Fig. 3c) and were not clearly distinguished from frog samples when considering phylogenetic diversity (Fig. 3a**, b**). Given that the effect of site was relatively minor and the microbial communities of frogs from Ponds 1 and 2 were largely similar, we combined these groups in further analysis.

**Fig. 3.**
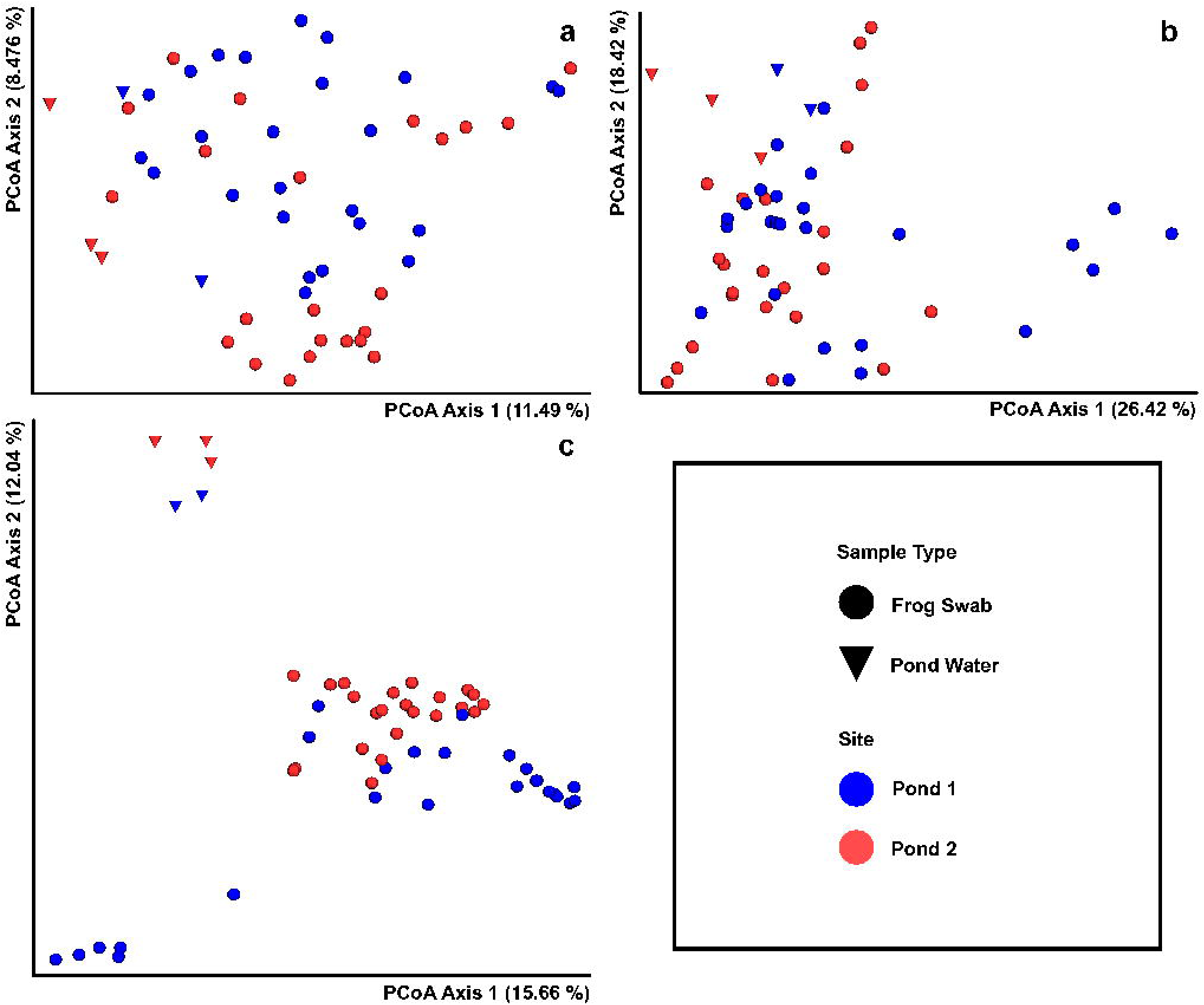
Principal coordinate analysis of β-diversity of spring frog skin and pond water microbiome samples. Principal coordinate analysis plots were created using Emperor from distance matrices calculated using a sampling depth of 10,000. Plots were limited to representing the two dimensions with the highest percent variation explained and were calculated for (**a)** Unweighted UniFrac distances, **(b)** Weighted UniFrac distances, and (**c)** Bray-Curtis distances

**Table 2.** Summary of *adonis* (PERMANOVA) models of β-diversity for microbial communities on frog skin during spring months and in pond water samples. Effects on variation due to sample type (frog skin, water), site (Pond 1, Pond 2) and year (2018, 2019) are considered. Significant results are marked with an asterisk

| Variables | Degrees of Freedom | Sums of Squares | Mean Squares | F.Model | R<sup>2</sup> | p |
| --- | --- | --- | --- | --- | --- | --- |
| **a) Unweighted UniFrac Distance** |  |  |  |  |  |  |
| Sample Type | 1 | 0.69 | 0.69 | 4.08 | 0.07 | 0.001* |
| Site | 1 | 0.48 | 0.48 | 2.85 | 0.05 | 0.001* |
| Year | 1 | 0.70 | 0.70 | 4.20 | 0.07 | 0.001* |
| Residuals |  | 8.23 | 0.17 |  | 0.81 |  |
| Total |  | 10.10 |  |  | 1.00 |  |
| **b) Weighted UniFrac Distance** |  |  |  |  |  |  |
| Sample Type | 1 | 0.66 | 0.66 | 6.86 | 0.11 | 0.001* |
| Site | 1 | 0.30 | 0.30 | 3.13 | 0.05 | 0.005* |
| Year | 1 | 0.44 | 0.44 | 4.54 | 0.07 | 0.001* |
| Residuals |  | 4.70 | 0.10 |  | 0.77 |  |
| Total |  | 6.10 |  |  | 1.00 |  |
| **c) Bray-Curtis Dissimilarity** |  |  |  |  |  |  |
| Sample Type | 1 | 1.83 | 1.83 | 7.71 | 0.12 | 0.001* |
| Site | 1 | 1.12 | 1.12 | 4.70 | 0.07 | 0.001* |
| Year | 1 | 1.28 | 1.28 | 5.39 | 0.08 | 0.001* |
| Residuals |  | 11.64 | 0.24 |  | 0.73 |  |
| Total |  | 15.87 |  |  | 1.00 |  |

### 3.3 Host Sex Does Not Affect Diversity and Structure of Microbial Communities

Due to sampling limitations, very few female frogs (n = 8) were included compared to the number of male frogs (n = 52). All female frogs were captured during 2018, and the majority (n = 6) were captured during spring, meaning that any sex-dependent microbiome characteristics might contribute to apparent seasonal differences. We found sex did not significantly affect microbiome diversity or structure. Comparing the α-diversity metrics of all male and female frogs using a t-test with Welch’s correction resulted in no significant difference for any metric observed (ASV richness: p = 0.3863; Shannon diversity index: p = 0.2717; Faith’s phylogenetic diversity: p = 0.4585). Additionally, sex was not a significant driver of community structure when assessed using *adonis* for the three β-diversity metrics used (Unweighted UniFrac: pseudo-F = 1.13, p = 0.241, R^2^ = 0.019; Weighted UniFrac: pseudo-F = 1.83, p = 0.097, R^2^ = 0.031; Bray-Curtis: pseudo-F = 1.30, p = 0.147, R^2^ = 0.022).

### 3.4 Season of Capture Influences the Structure of Microbial Communities

To assess the influence of season on the skin microbiome the frog samples were divided into groups based on capture date (Spring 2018, Summer 2018, Fall 2018 and Spring 2019). The microbial communities of frog skin from each seasonal group were similarly rich and even (Fig. 4). Of the three α-diversity metrics we considered, two differed significantly between seasonal sample groups (ANOVA: ASV richness, p < 0.0001; Faith’s phylogenetic diversity, p < 0.0025). In both cases the mean diversity of the Spring 2019 group was significantly higher than that of Spring 2018 and Summer 2018, while no other groups had significantly different means.

**Fig. 4.**
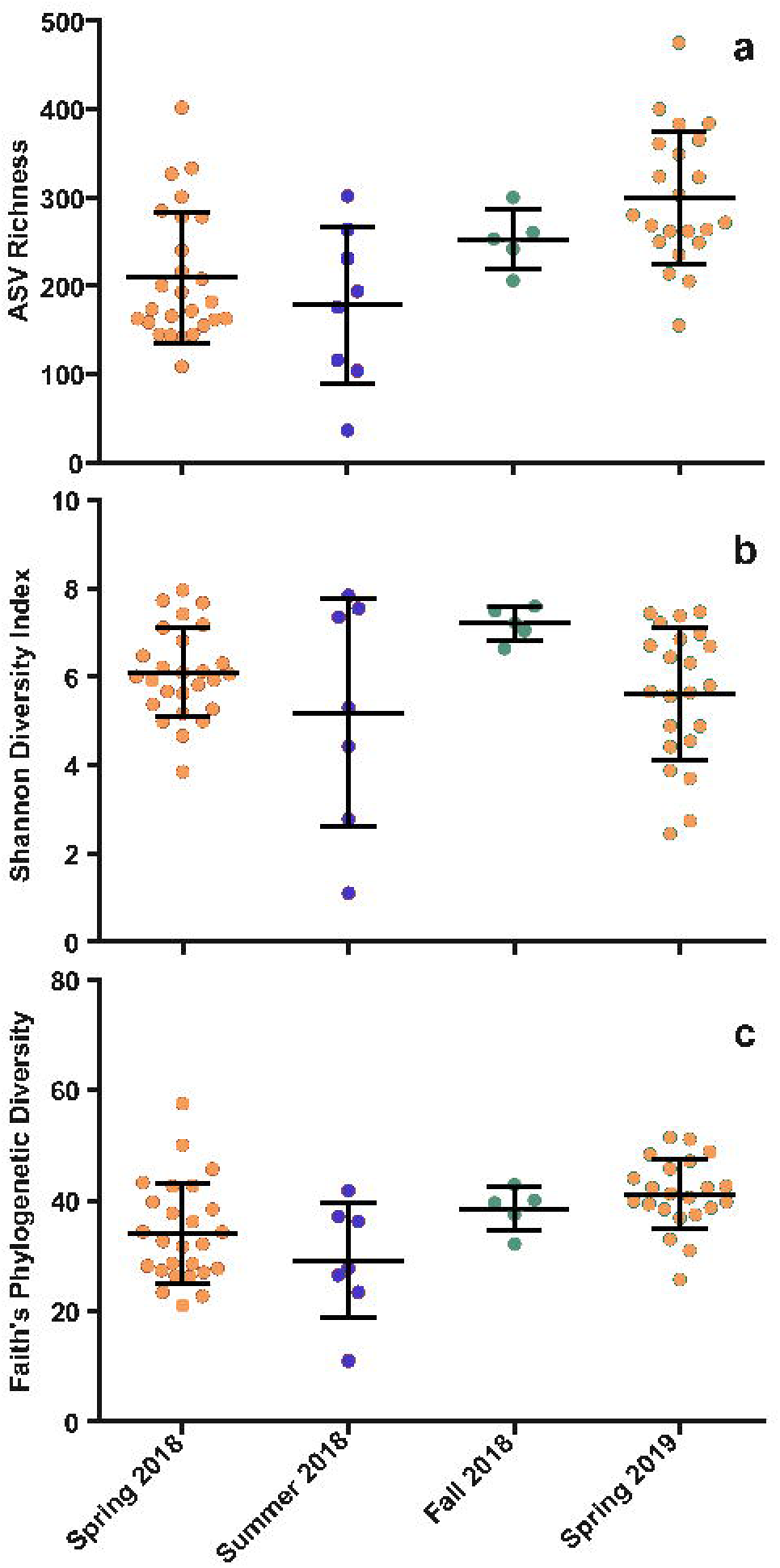
Comparison of seasonal wood frog skin microbiota α-diversity diversity metrics. Sample was α-diversity calculated using a sampling depth of 10,000. Mean seasonal value and standard deviation of each group is shown. Results are given for (**a)** ASV Richness, (**b)** Shannon’s Diversity Index and (**c)** Faith’s Phylogenetic Diversity

We found that community structure differed between seasonal groups, including observable trends at the highest taxonomic levels (Fig. 2). While the majority of ASVs present in any given sample were typically Proteobacteria, the abundance of ASVs belonging to other major phyla such as Bacteroidetes, Actinobacteria, Verrucomicrobia and Acidobacteria varied considerably (Fig. 2). Five samples from Spring 2019 were notable outliers, which had a unique community structure not observed in other samples. These frogs were all captured on the first day of sampling in Spring 2019 at Pond 1 and were the only samples where ASVs belonging to Actinobacteria made up more than 40% of the sequences detected, while those belonging to Proteobacteria were much less abundant (below 40%). We assessed the variation in relative abundance of the 5 major phyla between seasonal groups using one-way ANOVA. All groups exhibited some seasonal variation (Acidobacteria, p < 0.0001; Actinobacteria, p = 0.0138; Bacteroidetes, p = 0.0345; Proteobacteria, p = 0.0002; Verrucomicrobia, p = 0.0389). Acidobacteria was the only phylum showing a clear trend, with a significantly higher mean relative abundance (pairwise ANOVA: p < 0.0219) in Summer and Fall 2018 (4.7% and 6.4% respectively) than Spring 2018 and 2019 (1.9% and 0.62%, respectively).

Season was a significant source of variation in community structure when measured using Unweighted UniFrac (Table 3a, *adonis* pseudo-F = 4.36, p < 0.001, R^2^ = 0.13), Weighted UniFrac (Table 3b, *adonis* pseudo-F = 7.04, p < 0.001, R^2^ = 0.20) and Bray-Curtis (Table 3c, *adonis* pseudo-F = 6.17, p < 0.001, R^2^ = 0.18). We performed pairwise PERMANOVAs to determine which seasonal groups had significantly different community structure and every pairwise comparison revealed significant difference across all β-diversity metrics (PERMANOVA: p = 0.001), with the exception of the Summer and Fall 2018 frog skin microbiome groups which were not significantly different when considering Weighted UniFrac distance (PERMANOVA: pseudo-F = 2.60, p = 0.102). Summer and Fall 2018 frog skin microbiomes from were found to be significantly different when considering Unweighted UniFrac distance (PERMANOVA: pseudo-F = 1.69, p = 0.045) and Bray-Curtis dissimilarity (PERMANOVA: pseudo-F = 1.84, p = 0.045), though we noted that p values were near-threshold in both cases. This result was reflected in the Principle Coordinate Analysis results (Fig. 5), where skin microbiota from frogs collected during the summer and fall tended to cluster together, and appeared to form a cluster distinct from the spring samples when Unweighted UniFrac (Fig. 5a) and Bray-Curtis distances (Fig. 5c) were considered, although summer and fall samples were not clearly distinct from spring samples when visualizing Weighted Unifrac distances (Fig. 5b).

**Fig. 5.**
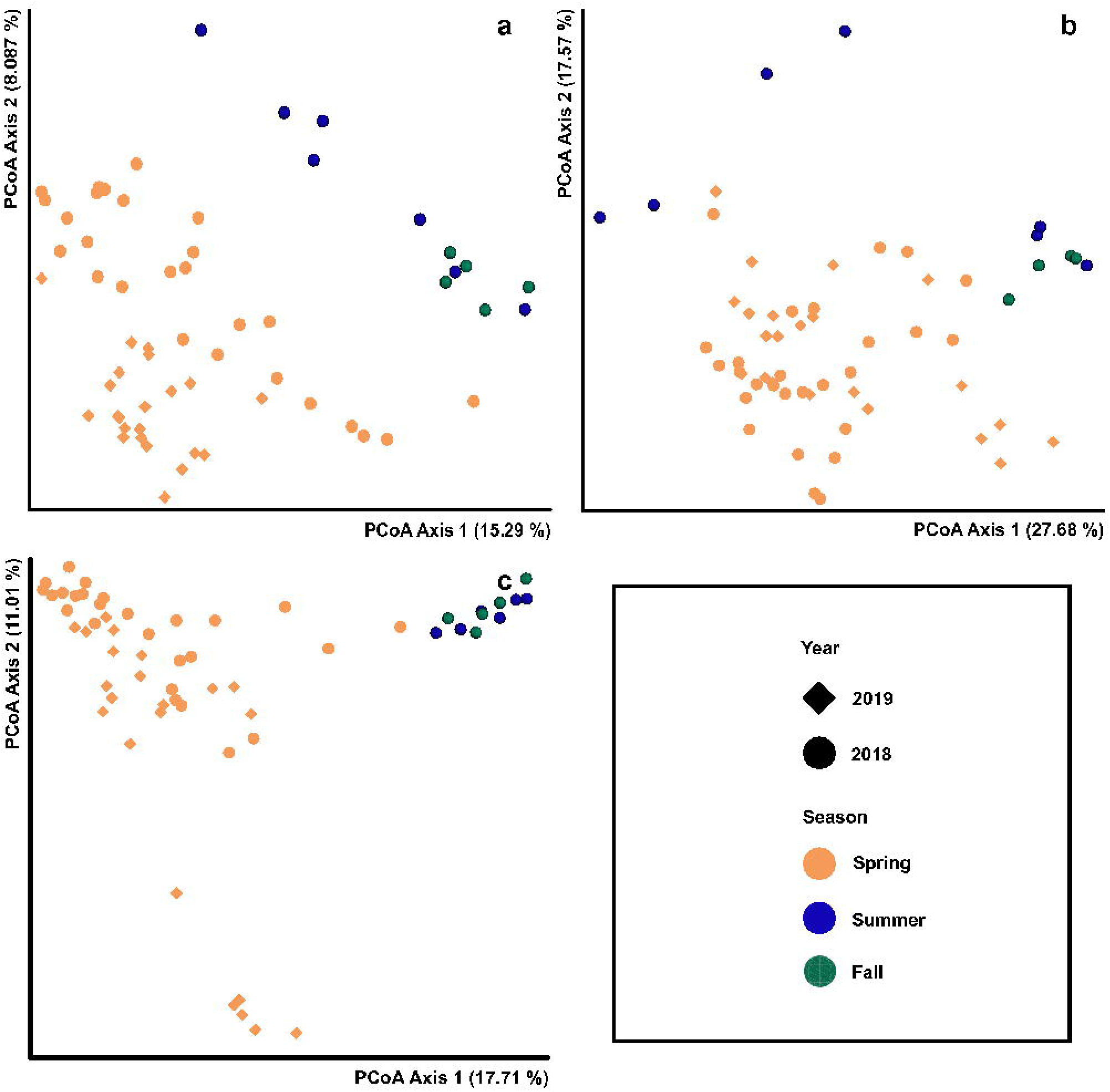
Principal coordinate analysis of β-diversity of frog skin microbiome samples. Principal coordinate analysis plots were created using Emperor from distance matrices calculated using a sampling depth of 10,000. Plots were limited to representing the two dimensions with the highest percent variation explained and were calculated for **(a)** Unweighted UniFrac distances, **(b)** Weighted UniFrac distances, and (**c)** Bray-Curtis distances

**Table 3.**
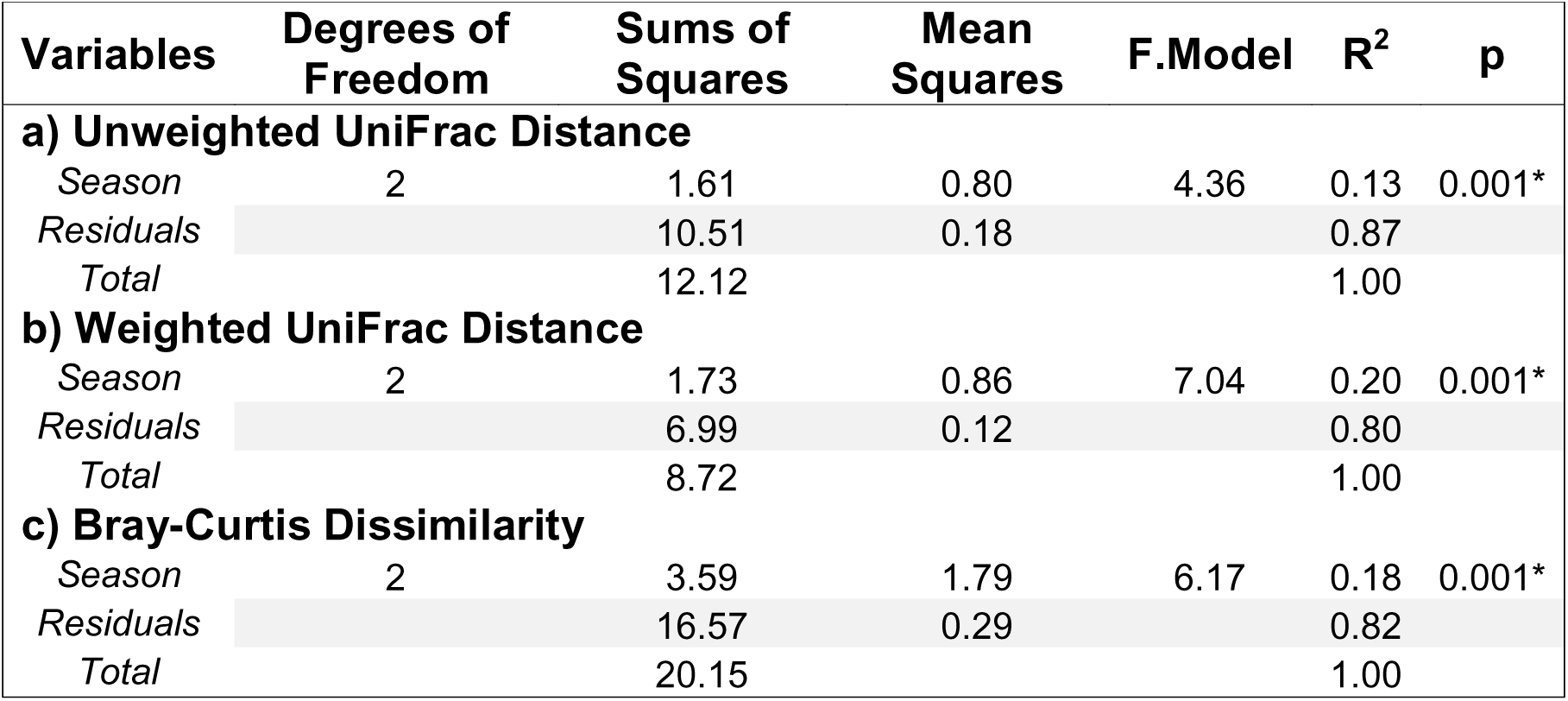
Summary of *adonis* (PERMANOVA) models of β-diversity for microbial communities on frog skin swab samples. Effects on variation due to season (spring, summer, fall) are considered. Significant results are marked with an asterisk

| Variables | Degrees of Freedom | Sums of Squares | Mean Squares | F.Model | R<sup>2</sup> | p |
| --- | --- | --- | --- | --- | --- | --- |
| **a) Unweighted UniFrac Distance** |  |  |  |  |  |  |
| Season | 2 | 1.61 | 0.80 | 4.36 | 0.13 | 0.001* |
| Residuals |  | 10.51 | 0.18 |  | 0.87 |  |
| Total |  | 12.12 |  |  | 1.00 |  |
| **b) Weighted UniFrac Distance** |  |  |  |  |  |  |
| Season | 2 | 1.73 | 0.86 | 7.04 | 0.20 | 0.001* |
| Residuals |  | 6.99 | 0.12 |  | 0.80 |  |
| Total |  | 8.72 |  |  | 1.00 |  |
| **c) Bray-Curtis Dissimilarity** |  |  |  |  |  |  |
| Season | 2 | 3.59 | 1.79 | 6.17 | 0.18 | 0.001* |
| Residuals |  | 16.57 | 0.29 |  | 0.82 |  |
| Total |  | 20.15 |  |  | 1.00 |  |

### 3.5 The Seasonal Core Microbiome

We identified core microbiota for all frog skin samples as a whole, as well as for each seasonal group separately. No ASV was present in ≥90% of all frog skin microbiome samples, so ASVs were collapsed at the genus and family level to consider a more inclusive core microbiome. A group of 7 genera were present in ≥90% of all frog samples, accounting for approximately 11% of ASVs on the average individual (Table 4). A group of 11 families were present in ≥90% of all frog samples, accounting for 56% of ASVs on the average individual (Table 5). We analyzed the seasonal variation in abundance of the nine core families with a mean relative abundance greater than 1% (Fig. 6), revealing six bacterial families with significantly different mean relative abundance across seasonal groups: Beijerinckaceae, Burkholderiaceae, Caulobacteraceae, Sphingobacteriaceae, Spirosomaceae and Xanthobacteracea (ANOVA: p < 0.05). In all cases but Sphingobacteriaceae, Summer and Fall 2018 abundances did not differ significantly. Similarly, Spring 2018 and 2019 significantly differed only in the abundance of Burkholderiaceae.

**Fig. 6.**
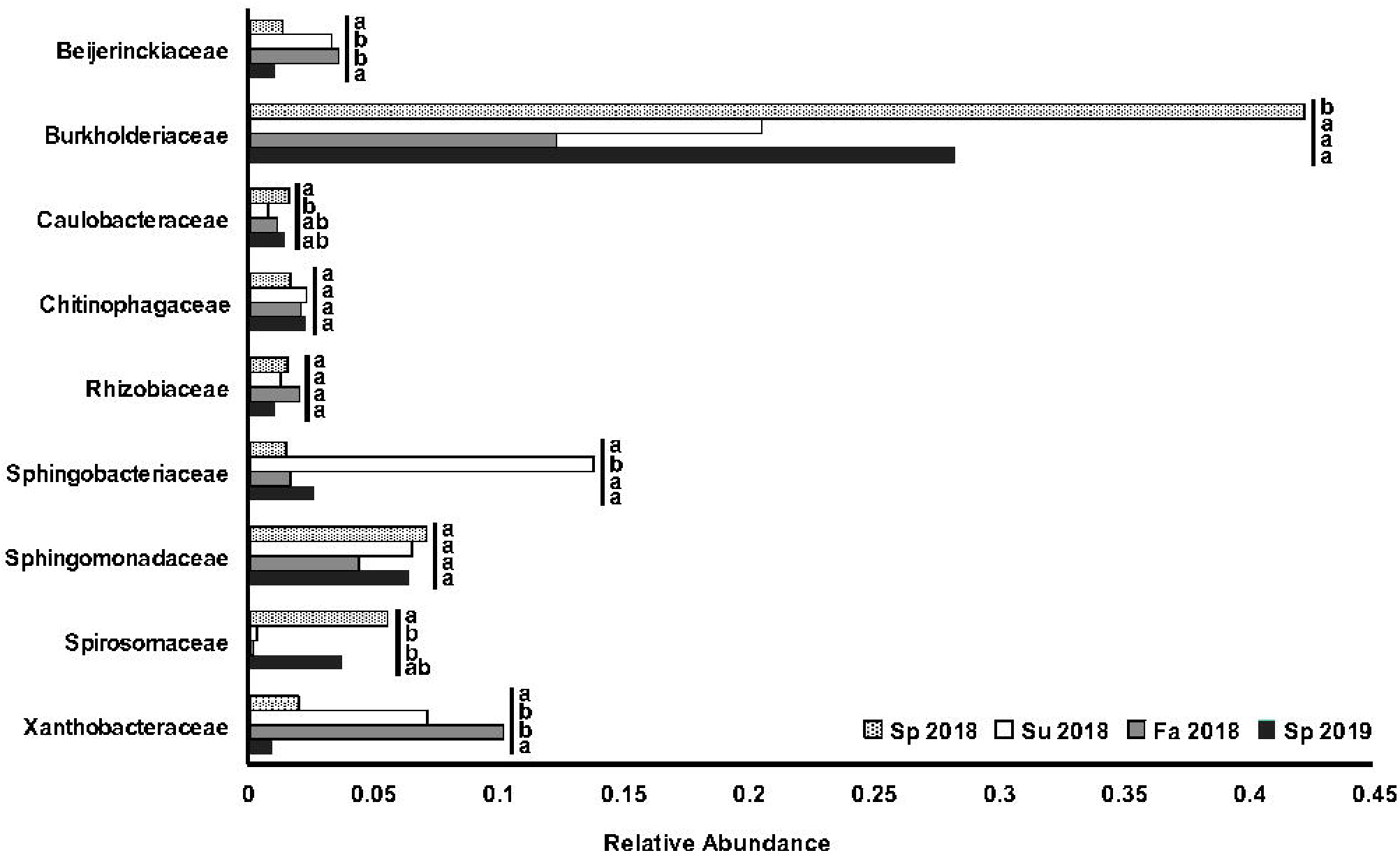
Relative abundance of core microbial families in wood frog skin microbiota. Families included were present in ≥ 90% of all frog skin samples and had a mean relative abundance ≥ 1%. Abundance of each family was compared between seasonal groups using pair-wise ANOVA, letters are used to indicate significant inter-seasonal variation for a given family. Seasons marked with the same letter do not significantly differ.

**Table 4.** Core microbiome genera and their mean relative abundance. Listed genera are present in ≥90% of all frog skin swab samples. If a microbial genus was not core to every seasonal group (“All”), seasonal groups for which the microbial genera was present in ≥90% of individual frog skin swab samples are listed. Wood frog skin swabs collected during different seasons are denoted in abbreviated form [spring (Sp), summer (Su) and fall (Fa) and corresponding year ((20)18 or 19)]

| Genera | Phylum | Mean Relative Abundance | Seasonally Core |
| --- | --- | --- | --- |
| Ferruginobacter | Bacteroidetes | 0.009 | Sp18, Sp19 |
| Uncultured<br>Chitinophagaceae | Bacteroidetes | 0.007 | Sp18, Su18, Sp19 |
| Methylobacterium | Proteobacteria | 0.006 | Su18, Fa18, Sp19 |
| Allo/Neo/Para/Rhizobium | Proteobacteria | 0.008 | All |
| Sphingomonas | Proteobacteria | 0.038 | All |
| Massilia | Proteobacteria | 0.019 | Sp18, Sp19 |
| Pseudomonas | Proteobacteria | 0.026 | Su18, Sp19 |

**Table 5.** Core microbiome families and their mean relative abundance. Listed families are present in ≥90% of all frog skin swab samples. If a microbial family was not core to every seasonal group (“All”), seasonal groups for which the microbial family was present in ≥90% of individual frog skin swab samples are listed. Wood frog skin swabs collected during different seasons are denoted in abbreviated form [spring (Sp), summer (Su) and fall (Fa) and corresponding year ((20)18 or 19)]

| Family | Phylum | Mean Relative Abundance | Seasonally Core |
| --- | --- | --- | --- |
| Chitinophagaceae | Bacteroidetes | 0.019 | All |
| Spirosomaceae | Bacteroidetes | 0.037 | Sp18, Sp19 |
| Sphingobacteriaceae | Bacteroidetes | 0.039 | Sp18, Su18, Sp19 |
| Acetobacteraceae | Proteobacteria | 0.009 | Su18, Fa18, Sp19 |
| Caulobacteraceae | Proteobacteria | 0.014 | All |
| Beijerinckiaceae | Proteobacteria | 0.014 | All |
| Rhizobiaceae | Proteobacteria | 0.013 | All |
| Xanthobacteraceae | Proteobacteria | 0.021 | All |
| Sphingomonadaceae | Proteobacteria | 0.062 | All |
| Burkholderiaceae | Proteobacteria | 0.329 | All |
| Xanthomonadaceae | Proteobacteria | 0.00401 | All |

When considering each seasonal group of skin associated communities separately, it was revealed that 25 of the 52 core genera and 16 of the 43 core families were considered core during only one season (Fig. 7). Only two genera were core to all seasonal groups (Fig. 7a), indicating that the number of core microbiota observed when considering all frogs together (7) was not reflective of the core genera present during each individual season. The Spring 2018 and Spring 2019 frog skin microbial communities had the most core genera in common (20), while Summer 2018 and Fall 2018 had the second highest level of overlap (7). Additionally, the frog skin microbiota from Spring 2018 and Spring 2019 each had a larger number of in-season core genera, with 20 in Spring 2018 and 37 in Spring 2019 as compared to 13 in Summer 2018 and 14 in Fall 2018. In comparison, a group of 8 families were core across all seasonal groups (Fig. 7b), but the consistent prevalence of these families was not always matched with a high relative abundance. We found that Burkholderiaceae was the only family to consistently have a mean relative abundance above 10% on wood frog skin, ranging from 13% in Fall 2018 to 44% in Spring 2018. Interestingly, there were 9 core families unique to the Fall 2018 wood frog skin group, while other seasonal groups had only 2-3 uniquely core families. Fall 2018 wood frog microbiomes also had the most core families overall at 26, followed by microbiota on wood frog skin in Spring 2019 with 25, Summer 2018 with 21, and Spring 2018 with 19. In all seasons except for Spring 2018, the core families present on wood frog skin were members of the phyla Actinobacteria, Bacteroidetes, Proteobacteria and Verrucomicrobia. The core families present on the skin of wood frogs in Spring 2018 lacked any members of Actinobacteria but included a single representative from the Gemmatimonadetes. In all cases the majority of core families present on wood frog skin were from the Proteobacteria, which ranged from 44% to 68% summed relative abundance. In general, community structure appeared to be more consistent for microbial families than for individual genera.

**Fig. 7.**
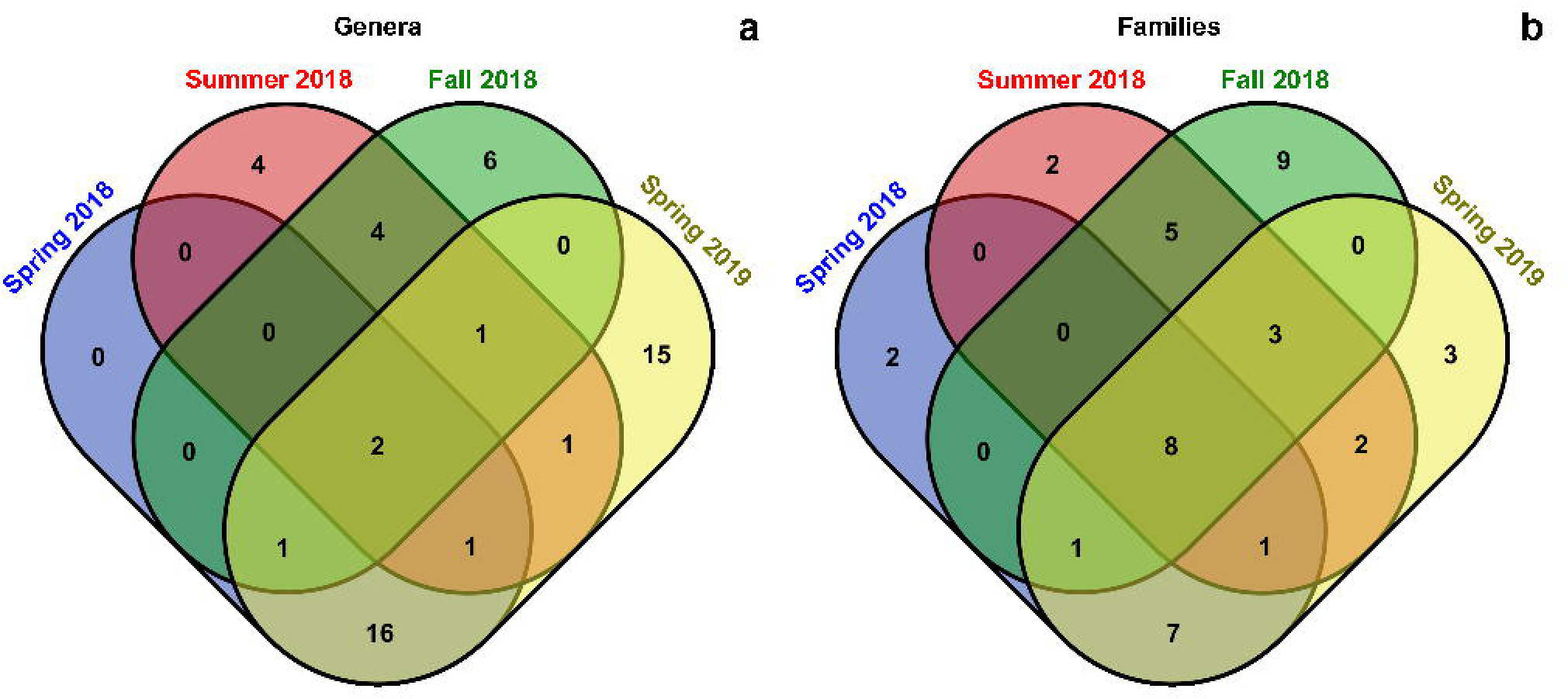
Overlap of core microbial families present on wood frog skin across seasons. Core taxa were defined as those found in 90% or more samples from a given season. Taxa were combined at the level of (**a)** genus and (**b)** family, omitting entries with ambiguous taxonomy

### 3.6 Putative Antifungal Taxa Are Present on the Skin of Wood Frogs Across Seasons

As recent studies have identified the importance of key bacterial species in protection of amphibians against *B. dendrobatidis* through the production of antifungal metabolites, we surveyed the wood frog microbiome for the presence of these putative antifungal bacterial species. Of the 37 bacterial genera found in the Antifungal Isolates Database that have isolated representatives demonstrating antifungal properties, 33 were present on *R. sylvatica* skin. The core genera from each seasonal group included putatively antifungal genera with several among the most prevalent genera overall, including *Pseudomonas*, *Massilia*, and *Allo/Neo/Para/Rhizobium*. Antifungal-associated genera varied in their relative abundances between wood frog skin microbiota samples, and mean relative abundances typically varied widely across seasons (Table 6).

**Table 6.** Putatively antifungal genera and their seasonal relative abundances. Genera which are found on *R. sylvatica* skin and have antifungal isolates listed in the Antifungal Isolates Database (Woodhams et al., 2015) are listed. Darker shading indicates greater relative abundance

| Genus | Mean Relative Abundance |  |  |  |
| --- | --- | --- | --- | --- |
|  | Spring 2018 | Summer 2018 | Fall 2018 | Spring 2019 |
| *Acinetobacter* | 0.008 | 0.004 | 0.000 | 0.000 |
| *Aeromonas* | 0.003 | 0.001 | 0.001 | 0.000 |
| *Allorhizobium-Neorhizobium-Pararhizobium-Rhizobium* | 0.008 | 0.006 | 0.014 | 0.007 |
| *Bacillus* | 0.000 | 0.001 | 0.002 | 0.000 |
| *Brevundimonas* | 0.010 | 0.001 | 0.001 | 0.006 |
| *Burkholderia-Caballeronia-Paraburkholderia* | 0.000 | 0.006 | 0.051 | 0.001 |
| *Chryseobacterium* | 0.015 | 0.001 | 0.000 | 0.016 |
| *Comamonas* | 0.000 | 0.001 | 0.001 | 0.000 |
| *Curtobacterium* | 0.004 | 0.013 | 0.003 | 0.162 |
| *Duganella* | 0.011 | 0.001 | 0.002 | 0.006 |
| *Dyella* | 0.000 | 0.001 | 0.010 | 0.000 |
| *Flavobacterium* | 0.015 | 0.000 | 0.008 | 0.029 |
| *Janthinobacterium* | 0.003 | 0.000 | 0.000 | 0.003 |
| *Lactococcus* | 0.000 | 0.000 | 0.000 | 0.000 |
| *Lysobacter* | 0.000 | 0.001 | 0.001 | 0.000 |
| *Massilia* | 0.030 | 0.002 | 0.004 | 0.018 |
| *Micrococcus* | 0.000 | 0.002 | 0.000 | 0.000 |
| *Novosphingobium* | 0.006 | 0.002 | 0.003 | 0.009 |
| *Paenibacillus* | 0.005 | 0.006 | 0.007 | 0.005 |
| *Pantoea* | 0.000 | 0.001 | 0.000 | 0.000 |
| *Pedobacter* | 0.007 | 0.003 | 0.003 | 0.019 |
| *Polaromonas* | 0.003 | 0.000 | 0.003 | 0.009 |
| *Pseudomonas* | 0.012 | 0.009 | 0.037 | 0.037 |
| *Pseudoxanthomonas* | 0.000 | 0.000 | 0.000 | 0.000 |
| *Serratia* | 0.000 | 0.000 | 0.000 | 0.000 |
| *Silvimonas* | 0.000 | 0.000 | 0.003 | 0.000 |
| *Sphingobacterium* | 0.000 | 0.001 | 0.000 | 0.000 |
| *Staphylococcus* | 0.001 | 0.004 | 0.017 | 0.000 |
| *Stenotrophomonas* | 0.000 | 0.001 | 0.006 | 0.001 |
| *Streptomyces* | 0.001 | 0.001 | 0.005 | 0.000 |
| *Undibacterium* | 0.007 | 0.000 | 0.000 | 0.009 |
| *Variovorax* | 0.005 | 0.006 | 0.007 | 0.005 |

## 4 Discussion

As we continue to improve our understanding of the skin microbiome and its role in maintaining the health of amphibians it is important to consider the inherent variability in microbial communities and the factors that drive this variability. While amphibians found in warmer regions may experience a relatively stable environment year-round, the majority of North America amphibians experience major seasonal fluctuations in environmental conditions. Changes in environmental conditions are known to influence the microbiome [4, 6, 7], potentially affecting the hosts ability to resist infection by a pathogen [18, 19]. In this study we have reported on the microbial community associated with *R. sylvatica* skin over the course of multiple seasons to observe the changes in community composition and structure. To provide perspective for these seasonal effects, we have compared them to the effects of host sex and vernal pool site of capture within a single season (spring). Lastly, we have highlighted members of the wood frog skin microbial community that are reported in the Antifungal Isolates Database to produce molecules with antifungal activity, to tentatively predict the impact of season on the potential ability of the skin microbiome to contribute to defense against fungal pathogens.

### 4.1 Microbial Community Structure and Core Taxa

We observed that the microbial community associated with *R. sylvatica* skin has much in common with those found on other frog species. ASVs belonging to Proteobacteria, Actinobacteria, Bacteroidetes, Firmicutes, Verrucomicrobia and Acidobacteria made up the vast majority of sequences observed from all samples, suggesting that these phyla dominate the microbiome. These phyla are commonly present on the skin of other frog species [3–5, 8] and all but Verrucomicrobia are abundant elements of the microbial communities associated with the related frog species *Rana pipiens* and *Rana catesbeiana* [1, 41]. While these phyla varied in abundance on the skin of individual wood frogs, there were very few cases in which any of the above bacterial phyla were found to be absent from the wood frog skin microbiota. Communities were much less consistent at finer taxonomic levels and among the hundreds of microbial genera observed, very few were prevalent enough to be considered core taxa. The most abundant of these core genera, *Sphingomonas* and *Pseudomonas*, are widespread in the environment at large and have been shown to be similarly abundant on the skin of other frogs [1, 6, 41].

Many of the core taxa associated with the skin microbiome are also known contaminants of commercial DNA extraction kits [42]. To better understand the extent to which microbial contaminants introduced during the extraction process contributed to the microbial communities observed in our samples, we looked for ASVs present in both the samples and the process controls. While the majority of our frog samples (n = 43) had a low abundance (>5% relative abundance) of ASVs which were found in the process controls, a group of seven frog skin samples from Summer 2018 and Spring 2019 had very high (>50%) relative abundance of ASVs found in the process controls, which seems to indicate a high level of process contamination in these samples. Unexpectedly, all water samples had a very low abundance of ASVs found in the process control (0.07% average relative abundance) and completely lacked the *Curtobacterium* ASV which was highly abundant in process blanks, field blanks and frog samples. This suggests that potential contamination from reagents was not universal, or that many of the ASVs detected in the process controls were also naturally present on the frogs and surrounding environment. Additionally, samples with low numbers of detected amplicons did not have a proportionally higher relative abundance of ASVs found in the process controls, as would be expected of a failed skin swab which did not capture frog skin microbiota. Overall, while contamination from reagents and exposure to the laboratory environment was impossible to avoid, it does not appear to contribute to the observed trends in community structure.

### 4.2 Effect of Vernal Pool Site and Host Sex on the Wood Frog Skin Microbiome

While *R. sylvatica* are generally terrestrial and solitary, wood frogs converge on vernal pools during the spring thaw to seek mates and reproduce [23]. In this study the two temporary ponds from which frogs were captured during the spring served as the only truly distinct sampling sites, since the surrounding area was fairly uniform mixed woodland. As *R. sylvatica* are known to venture as far as 1 km from their breeding pond and the ponds sampled are ∼200 m apart it is unlikely that they harbor genetically distinct populations [43], and therefore any variation in the skin-associated microbial community is better attributed to the environmental conditions of the site. This is an important distinction, as it is not well understood to what degree host phylogenetics and environment affect the microbiome of amphibians, and examples exist which emphasize the role of both factors [44–46]. Pond of origin was found to explain a small, but significant amount of the variation observed in the microbial communities on frogs captured during the spring. This effect was less pronounced than the variation between frogs captured in spring of 2018 and 2019 however, and the largest differences were observed between frog swabs and water samples. This was expected, as previous studies have established that amphibians have communities of skin-associated microbiota distinct from their environment [1, 41, 47]. There was no clear trend linking frog skin microbiomes to the microbiota found in the associated pond water. There was no significant difference in ASV richness, evenness or phylogenetic diversity between the skin-associated microbial communities in Pond 1 and Pond 2. Despite water samples from Pond 2 having a lower mean ASV richness than those of Pond 1, frogs from Pond 2 had more ASVs on average, suggesting that seeding of microbes from the water was not a major driver of skin microbiome diversity. Frogs from both ponds hosted bacterial phyla that were uncommon, or not present, in the pond water and exhibited more diverse and even communities, while water samples were almost entirely populated by Proteobacteria, Bacteroidetes and Actinobacteria. Given the increased abundance of many of these phyla on frogs captured during the summer and fall, is seems they must either be stable members of the microbiome or are seeded from rich microbial communities found in the soil and leaf litter of the surrounding forest. It is unclear whether seeding from soil environments might occur while buried during winter hibernation, and a study of the microbial communities present in frog hibernacula, although challenging, would be an interesting avenue of future research.

The effect of frog sex on the microbiome was also considered, as relationships between sex and skin microbiota have been observed in humans [48] and other vertebrates [49], but the effect of sex has not been well studied in amphibians. We found sex had no significant effect on structure or diversity of the microbial community. The few previous studies considering the effect of sex in amphibians failed to find significant differences between males and females [21, 44], and our work, although limited by the low number of female frogs, corroborates these findings.

### 4.3 Effect of Season on the Wood Frog Skin Microbiome

Season was also associated with significant variation in the structure and composition of the wood frog skin microbiomes studied. The effect of season was more pronounced than the effects of site, year or sample type observed in the spring samples, and was evidenced by shifts in the abundance of major phyla on wood frog skin. Most notably, relative abundance of Acidobacteria was significantly higher among frogs captured in the summer and fall than those captured during spring. Given the particularly high abundance of Acidobacteria in soils [50] it is not surprising that members of this phylum would be highly abundant on frogs active in soil and leaf litter. The seasonality to Acidobacteria on frog skin matches a proposal that Acidobacteria are transiently associated with the human skin microbiome [51]. While the composition of the frog skin microbiome varied, average diversity of individual *R. sylvatica* microbiomes remained fairly constant across seasons. When considering ASV richness and phylogenetic diversity the Spring 2019 wood frog skin microbiota group was determined to have a mean diversity significantly different from Spring 2018 and Summer 2018. However, when considering the Shannon Diversity Index, wood frog skin microbiomes sampled during Spring 2019 falls well within the range of these groups. This suggests that the Spring 2019 wood frog microbial communities were not as even, and a larger proportion of uncommon ASVs were contributing to their diversity. This is likely at least partially the result of the Spring 2019 samples undergoing an additional sequencing replicate. The resultant greater sequencing depth would increase the number of rare ASVs [52], and contribute to increased observed diversity. Rarefaction prior to diversity analyses was conducted to mitigate this issue, but it is not a perfect method [53].

Due to the much higher number of wood frog microbiome samples collected during the spring months (75% of total frog swab samples), any analysis considering overall prevalence of taxa was heavily biased toward taxa which were common during the spring. To better represent the microbial communities present on wood frog skin during summer and fall samples, core taxa were considered for each seasonal group individually and overlaps in seasonal groups’ core taxa determined. As core taxa represent the microbes which are most commonly found in the frog skin environment and often represent key members of the microbial community [39], common core taxa should reflect similar community dynamics. The skin of *R. sylvatica* hosted only a small number of core microbiota, particularly at lower taxonomic levels. Several of the core taxa are known to be core to other frog skin communities, *Pseudomonas* being one of the most commonly represented [6, 38, 41], but the majority of core taxa appeared to be fairly unique. Additionally, many of the most prevalent taxa experienced significant changes in abundance between seasons. The families Beijerinckiaceae and Xanthobacteraceae were significantly more abundant in summer and fall, while Sphingobacteriaceae greatly increased in abundance during the summer only. The variability in the core taxa observed on *R. sylvatica* skin suggests that the skin microbiome is a highly dynamic environment, where seasonal factors can re-shape the core structure and few ‘microbes are suited to inhabit the skin year-round.

### 4.4 Seasonal Representation of Putatively Anti-fungal Microbes

Like many who study the amphibian microbiome, we aimed to improve our understanding of the trends which contribute to resistance to major pathogens. Our analysis focused on members of the frog skin microbial community that protect against *B. dendrobatidis*. Notably, we observed members of the genus *Janthinobacterium* on *R. sylvatica* skin, however it was not determined whether the ASV detected belonged to the protective species *Janthinobacterium lividum* [10, 17]. *Janthinobacterium* were most abundant in Spring 2018 and 2019, were present on only one Summer 2018 frog and were entirely absent in Fall 2018. Additionally, members of the family Sphingobacteriaceae were found on frogs in all seasons. This is notable as presence of Sphingobacteriaceae was a predictor of successful recovery from *B. dendrobatidis* infection in other frog species [28]. Several other core taxa present on wood frog skin have known isolates which inhibit *B. dendrobatidis* listed in the Antifungal Isolates Database of amphibian skin-associated bacteria [40]. Burkholderiaceae and Xanthomonadaceae, in particular, have many genera with anti- *B. dendrobatidis* isolated members. Among the core genera, *Pseudomonas* and *Rhizobium* have the strongest evidence for *B. dendrobatidis* inhibition *in vitro* [40]. Aside from a spike in Burkholderiaceae abundance in Spring 2018, putatively anti-*B. dendrobatidis* taxa were not associated with any season in particular, and most were present at low abundance throughout the year. While these findings do not confirm that the bacterial taxa observed on *R. sylvatica* have antifungal activity, the organisms in our dataset associated with these groups are of interest as putatively anti-*B. dendrobatidis*. Further investigation is required to determine whether the specific microbial strains present on *R. sylvatica* possess anti-pathogen qualities to better understand the functional significance of seasonal variation in the skin microbiome and its contribution to defense against pathogens.

## 5 Conclusions

Our results indicate that season has a significant effect on the structure of the North American wood frog skin microbiome and has a proportionally greater effect than spring breeding pond association. Frogs captured during summer and fall were the most similar in terms of β-diversity distances and could be distinguished from spring frogs by their increased abundance of Acidobacteria, as well as other soil-associated bacterial families. It remains unclear whether the shift towards increased abundance of soil-associated bacteria on frog skin in the summer and fall is a result of transient colonization from frequent exposure, or a stable equilibrium shift in the community. Skin-associated microbial communities had consistent structural similarities at the highest taxonomic levels but displayed a high degree of diversity at finer levels, and the few core genera identified were not a dominant component of the community. Frogs captured during all seasons were host to microbes with putative anti- *B. dendrobatidis* activity, and seasonal shifts did not seem to affect the overall pool of potentially protective taxa. *R. sylvatica* is a widespread species and further study of populations from varied environments (Boreal shield, montane forest, etc.) could reveal related trends. While the effect of season has been briefly explored in other temperate frog species [4], this study provides insight into the seasonality of skin microbiome structure on amphibian species inhabiting northern environments and establishes foundational knowledge for further study of species which experience dramatic shifts in habitat and behavior between seasons.

## Acknowledgements

The authors thank Nicole Wang for the generous contribution of a trained taxonomic classifier for 16S rRNA gene sequences and Maxwell P. Bui-Marinos, Joseph F.A. Varga and Nathanael B. J. Harper for their technical assistance in collecting frog skin swabs.

## Author Contributions

AJD, LAH and BAK conceived the study; AJD and BAK performed field sampling; AJD performed the experiments and analyzed the data; AJD, LAH and BAK wrote and critically revised the manuscript.

## Data Availability

16S rRNA gene amplicon sequence data for skin microbiome samples are deposited in the NCBI Sequence Read Archive (Bioproject PRJNA603391).

## Compliance with Ethical Standards

### Ethics Statement

All applicable international, national, and/or institutional guidelines for the care and use of animals were followed. All procedures performed in studies involving animals were in accordance with the ethical standards of the institution at which the studies were conducted (University of Waterloo Animal Care Committee and the Canadian Council on Animal Care, Animal Utilization Projects #30008 and #40721; and animals captured under the Ontario Ministry of Natural Resources and Forestry Wildlife Scientific Collectors Authorization Permits #1088586 and #1092603 issued to Dr. B.A. Katzenback). This article does not contain any studies with human participants performed by any of the authors.

## References

1. McKenzie VJ, Bowers RM, Fierer N, et al (2012) Co-habiting amphibian species harbor unique skin bacterial communities in wild populations. ISME J 6:588–596. https://doi.org/10.1038/ismej.2011.129

2. Colombo BM, Scalvenzi T, Benlamara S, Pollet N (2015) Microbiota and mucosal immunity in amphibians. Front. Immunol. 6:1–15. https://doi.org/10.3389/fimmu.2015.00111

3. Walke JB, Becker MH, Hughey MC, et al (2015) Most of the dominant members of amphibian skin bacterial communities can be readily cultured. Appl. Environ. Microbiol. 81:6589–6600. https://doi.org/10.1128/AEM.01486-15

4. Longo A V., Savage AE, Hewson I, Zamudio KR (2015) Seasonal and ontogenetic variation of skin microbial communities and relationships to natural disease dynamics in declining amphibians. R. Soc. Open. Sci. 2:140377. https://doi.org/10.1098/rsos.140377

5. Belden LK, Hughey MC, Rebollar EA, et al (2015) Panamanian frog species host unique skin bacterial communities. Front. Microbiol. 6:1171. https://doi.org/10.3389/fmicb.2015.01171

6. Ellison S, Rovito S, Parra-Olea G, et al (2018) The Influence of Habitat and Phylogeny on the Skin Microbiome of Amphibians in Guatemala and Mexico. Microb. Ecol. 1–11. https://doi.org/10.1007/s00248-018-1288-8

7. Costa S, Lopes I, Proença DN, et al (2016) Diversity of cutaneous microbiome of *Pelophylax perezi* populations inhabiting different environments. Sci. Total Environ. 572:995–1004. https://doi.org/10.1016/j.scitotenv.2016.07.230

8. Varga JFA, Bui-Marinos MP, Katzenback BA (2019) Frog skin innate immune defences: Sensing and surviving pathogens. Front. Immunol. 10:3128. https://doi.org/10.3389/fimmu.2018.03128

9. Woodhams DC, Vredenburg VT, Simon MA, et al (2007) Symbiotic bacteria contribute to innate immune defenses of the threatened mountain yellow-legged frog, *Rana muscosa*. Biol. Conserv. 138:390–398. https://doi.org/10.1016/j.biocon.2007.05.004

10. Harris RN, Brucker RM, Walke JB, et al (2009) Skin microbes on frogs prevent morbidity and mortality caused by a lethal skin fungus. ISME J 3:818–824. https://doi.org/10.1038/ismej.2009.27

11. Brucker RM, Baylor CM, Walters RL, et al (2008) The identification of 2,4-diacetylphloroglucinol as an antifungal metabolite produced by cutaneous bacteria of the salamander *Plethodon cinereus*. J Chem. Ecol. 34:39–43. https://doi.org/10.1007/s10886-007-9352-8

12. Lauer A, Simon MA, Banning JL, et al (2007) Common Cutaneous Bacteria from the Eastern Red-Backed Salamander Can Inhibit Pathogenic Fungi. Copeia 2007:630–640. https://doi.org/10.1643/0045-8511(2007)2007[630:CCBFTE]2.0.CO;2

13. Lauer A, Simon MA, Banning JL, et al (2008) Diversity of cutaneous bacteria with antifungal activity isolated from female four-toed salamanders. ISME J 2:145–157. https://doi.org/10.1038/ismej.2007.110

14. Daszak P, Cunningham AA, Hyatt AD (2003) Infectious disease and amphibian population declines. Divers. Distrib. 9:141–150. https://doi.org/10.1046/j.1472-4642.2003.00016.x

15. Kilpatrick AM, Briggs CJ, Daszak P (2010) The ecology and impact of chytridiomycosis: an emerging disease of amphibians. Trends Ecol. Evol. 25:109–118. https://doi.org/10.1016/j.tree.2009.07.011

16. Harris RN, Lauer A, Simon MA, et al (2009) Addition of antifungal skin bacteria to salamanders ameliorates the effects of chytridiomycosis. Dis. Aquat. Organ. 83:11–16. https://doi.org/10.3354/dao02004

17. Kueneman JG, Woodhams DC, Harris R, et al (2016) Probiotic treatment restores protection against lethal fungal infection lost during amphibian captivity. Proc. R. Soc. B. Biol. Sci. 283:20161553. https://doi.org/10.1098/rspb.2016.1553

18. Muletz-Wolz CR, Almario JG, Barnett SE, et al (2017) Inhibition of fungal pathogens across genotypes and temperatures by amphibian skin bacteria. Front. Microbiol. 8:1551. https://doi.org/10.3389/fmicb.2017.01551

19. Ellison S, Knapp RA, Sparagon W, et al (2019) Reduced skin bacterial diversity correlates with increased pathogen infection intensity in an endangered amphibian host. Mol. Ecol. 28:127–140. https://doi.org/10.1111/mec.14964

20. Walke JB, Becker MH, Loftus SC, et al (2015) Community structure and function of amphibian skin microbes: An experiment with bullfrogs exposed to a chytrid fungus. PLoS One 10:e0139848. https://doi.org/10.1371/journal.pone.0139848

21. Campbell LJ, Garner TWJ, Hopkins K, et al (2019) Outbreaks of an Emerging Viral Disease Covary With Differences in the Composition of the Skin Microbiome of a Wild United Kingdom Amphibian. Front. Microbiol. 10:1245. https://doi.org/10.3389/fmicb.2019.01245

22. Berven KA, Gill DE (1983) Interpreting geographic variation in life-history traits. Integr. Comp. Biol. 23:85–97. https://doi.org/10.1093/icb/23.1.85

23. Berven KA (1990) Factors affecting population fluctuations in larval and adult stages of the wood frog (*Rana sylvatica*). Ecology 71:1599–1608. https://doi.org/10.2307/1938295

24. Matutte B, Storey KB, Knoop FC, Conlon JM (2000) Induction of synthesis of an antimicrobial peptide in the skin of the freeze-tolerant frog, *Rana sylvatica*, in response to environmental stimuli. FEBS Lett. 483:135–138. https://doi.org/10.1016/S0014-5793(00)02102-5

25. Storey KB, Storey JSKM (1992) Natural Freeze Tolerance In Ectothermic Vertebrates. Annu. Rev. Physiol. 54:619–637. https://doi.org/10.1146/annurev.physiol.54.1.619

26. Gahl MK, Longcore JE, Houlahan JE (2012) Varying Responses of Northeastern North American Amphibians to the Chytrid Pathogen *Batrachochytrium dendrobatidis*. Conserv. Biol. 26:135–141. https://doi.org/10.1111/j.1523-1739.2011.01801.x

27. Forzán MJ, Jones KM, Ariel E, et al (2017) Pathogenesis of Frog Virus 3 (Ranavirus, Iridoviridae) Infection in Wood Frogs (Rana sylvatica). Vet. Pathol. 54:531–548. https://doi.org/10.1177/0300985816684929

28. Becker MH, Walke JB, Cikanek S, et al (2015) Composition of symbiotic bacteria predicts survival in Panamanian golden frogs infected with a lethal fungus. Proc. R. Soc. B. Biol. Sci. 282:20142881–20142881. https://doi.org/10.1098/rspb.2014.2881

29. Rebollar EA, Gutiérrez-Preciado A, Noecker C, et al (2018) The skin microbiome of the neotropical frog *Craugastor fitzingeri*: Inferring potential bacterial-host-pathogen interactions from metagenomic data. Front. Microbiol. 9:466. https://doi.org/10.3389/fmicb.2018.00466

30. Walters W, Hyde ER, Berg-Lyons D, et al (2016) Improved Bacterial 16S rRNA Gene (V4 and V4-5) and Fungal Internal Transcribed Spacer Marker Gene Primers for Microbial Community Surveys. mSystems 1:e00009–15. https://doi.org/10.1128/mSystems.00009-15

31. Apprill A, Mcnally S, Parsons R, Weber L (2015) Minor revision to V4 region SSU rRNA 806R gene primer greatly increases detection of SAR11 bacterioplankton. Aquat. Microb. Ecol. 75:129–137. https://doi.org/10.3354/ame01753

32. Parada AE, Needham DM, Fuhrman JA (2016) Every base matters: Assessing small subunit rRNA primers for marine microbiomes with mock communities, time series and global field samples. Environ. Microbiol. 18:1403–1414. https://doi.org/10.1111/1462-2920.13023

33. Bolyen E, Rideout JR, Dillon MR, et al (2019) Reproducible, interactive, scalable and extensible microbiome data science using QIIME 2. Nat. Biotechnol. 37:852–857. https://doi.org/10.1038/s41587-019-0209-9

34. Callahan BJ, McMurdie PJ, Rosen MJ, et al (2016) DADA2: High-resolution sample inference from Illumina amplicon data. Nat. Methods. 13:581–583. https://doi.org/10.1038/nmeth.3869

35. Yilmaz P, Parfrey LW, Yarza P, et al (2014) The SILVA and “all-species Living Tree Project (LTP)” taxonomic frameworks. Nucleic Acids Res. 42:D643–D648. https://doi.org/10.1093/nar/gkt1209

36. Katoh, K., Misawa, K., Kuma, K., Miyata T (2002) MAFFT: a novel method for rapid multiple sequence alignment based on fast Fourier transform. Nucleic Acids Res. 30:3059–3066. https://doi.org/10.1093/nar/gkf436

37. Stamatakis A (2014) RAxML version 8: A tool for phylogenetic analysis and post-analysis of large phylogenies. Bioinformatics 30:1312–1313. https://doi.org/10.1093/bioinformatics/btu033

38. Medina D, Hughey MC, Becker MH, et al (2017) Variation in Metabolite Profiles of Amphibian Skin Bacterial Communities Across Elevations in the Neotropics. Microb. Ecol. 74:227–238. https://doi.org/10.1007/s00248-017-0933-y

39. Shade A, Handelsman J (2012) Beyond the Venn diagram: The hunt for a core microbiome. Environ. Microbiol. 14:4–12. https://doi.org/10.1111/j.1462-2920.2011.02585.x

40. Woodhams DC, Alford RA, Antwis RE, et al (2015) Antifungal isolates database of amphibian skin-associated bacteria and function against emerging fungal pathogens. Ecology 96:595–595. https://doi.org/10.1890/14-1837.1

41. Walke JB, Becker MH, Loftus SC, et al (2014) Amphibian skin may select for rare environmental microbes. ISME J 8:2207–2217. https://doi.org/10.1038/ismej.2014.77

42. Salter SJ, Cox MJ, Turek EM, et al (2014) Reagent and laboratory contamination can critically impact sequence-based microbiome analyses. BMC Biol. 12:87. https://doi.org/10.1186/s12915-014-0087-z

43. Berven KA, Grudzien TA (1990) Dispersal in the Wood Frog (*Rana sylvatica*): Implications for Genetic Population Structure. Evolution (N Y) 44:2047–2056. https://doi.org/10.2307/2409614

44. Prado-Irwin SR, Bird AK, Zink AG, Vredenburg VT (2017) Intraspecific Variation in the Skin-Associated Microbiome of a Terrestrial Salamander. Microb. Ecol. 74:745–756. https://doi.org/10.1007/s00248-017-0986-y

45. Muletz Wolz CR, Yarwood SA, Campbell Grant EH, et al (2018) Effects of host species and environment on the skin microbiome of Plethodontid salamanders. J. Anim. Ecol. 87:341–353. https://doi.org/10.1111/1365-2656.12726

46. Bletz MC, Archer H, Harris RN, et al (2017) Host ecology rather than host phylogeny drives amphibian skin microbial community structure in the biodiversity hotspot of Madagascar. Front. Microbiol. 8:1530. https://doi.org/10.3389/fmicb.2017.01530

47. Albecker MA, Belden LK, McCoy MW (2018) Comparative Analysis of Anuran Amphibian Skin Microbiomes Across Inland and Coastal Wetlands. Microb. Ecol. 78:348. https://doi.org/10.1007/s00248-018-1295-9

48. Fierer N, Hamady M, Lauber CL, Knight R (2008) The influence of sex, handedness, and washing on the diversity of hand surface bacteria. Proc. Natl. Acad. Sci. 105:17994–17999. https://doi.org/10.1073/pnas.0807920105

49. Saag P, Tilgar V, Mänd R, et al (2011) Plumage Bacterial Assemblages in a Breeding Wild Passerine: Relationships with Ecological Factors and Body Condition. Microb. Ecol. 61:740–749. https://doi.org/10.1007/s00248-010-9789-0

50. Kielak AM, Barreto CC, Kowalchuk GA, et al (2016) The ecology of Acidobacteria: Moving beyond genes and genomes. Front. Microbiol. 7:744 https://doi.org/10.3389/fmicb.2016.00744

51. Grönroos M, Parajuli A, Laitinen OH, et al (2019) Short-term direct contact with soil and plant materials leads to an immediate increase in diversity of skin microbiota. MicrobiologyOpen 8:e00645. https://doi.org/10.1002/mbo3.645

52. Zaheer R, Noyes N, Ortega Polo R, et al (2018) Impact of sequencing depth on the characterization of the microbiome and resistome. Sci. Rep. 8:5890. https://doi.org/10.1038/s41598-018-24280-8

53. McMurdie PJ, Holmes S (2014) Waste Not, Want Not: Why Rarefying Microbiome Data Is Inadmissible. PLoS Comput. Biol. 10:e1003531. https://doi.org/10.1371/journal.pcbi.1003531

