## Supplemental Table 1 for "Composition of the North American wood frog (*Rana sylvatica*) skin microbiome and seasonal variation in community structure"

**Supplementary Table 1** Bacterial taxa detected in procedural negative controls and their mean relative abundance. Three negative controls were produced by following the outlined DNA extraction and sequencing procedures without including any sample. Mean relative abundance values are given for genera where possible, or otherwise for most specific taxa known

| **Taxa** | **Mean Relative Abundance** |
| --- | --- |
| Actinobacteria;Actinobacteria;Micrococcales;Microbacteriaceae;Curtobacterium | 0.4293 |
| Firmicutes;Bacilli;Lactobacillales;Lactobacillaceae;Lactobacillus | 0.1573 |
| Proteobacteria;Alphaproteobacteria;Rhizobiales;Xanthobacteraceae | 0.0909 |
| Tenericutes;Mollicutes;Acholeplasmatales;Acholeplasmataceae;Acholeplasma | 0.0772 |
| Proteobacteria;Gammaproteobacteria;Betaproteobacteriales;Burkholderiaceae | 0.0518 |
| Bacteroidetes;Bacteroidia;Bacteroidales;Bacteroidaceae;Bacteroides | 0.0339 |
| Verrucomicrobia;Verrucomicrobiae;Verrucomicrobiales;Rubritaleaceae;Luteolibacter | 0.0202 |
| Bacteroidetes;Bacteroidia;Bacteroidales;Dysgonomonadaceae;Dysgonomonas | 0.0197 |
| Proteobacteria;Gammaproteobacteria;Enterobacteriales;Enterobacteriaceae;Escherichia-Shigella | 0.0156 |
| Chloroflexi;Gitt-GS-136 | 0.0122 |
| Firmicutes;Clostridia;Clostridiales;Clostridiaceae 1;Proteiniclasticum | 0.0116 |
| Actinobacteria;Actinobacteria;Micrococcales;Micrococcaceae;Paenarthrobacter | 0.0071 |
| Proteobacteria;Gammaproteobacteria;Betaproteobacteriales;Burkholderiaceae;Ralstonia | 0.0063 |
| Proteobacteria;Alphaproteobacteria;Rhizobiales;Rhizobiaceae;Mesorhizobium | 0.0059 |
| Bacteroidetes;Bacteroidia;Flavobacteriales;Flavobacteriaceae;Flavobacterium | 0.0051 |
| Proteobacteria;Gammaproteobacteria;Xanthomonadales;Xanthomonadaceae;Lysobacter | 0.0045 |
| Actinobacteria;Actinobacteria;Micrococcales;Micrococcaceae | 0.0039 |
| Proteobacteria;Gammaproteobacteria;Pseudomonadales;Pseudomonadaceae;Pseudomonas | 0.0034 |
| Proteobacteria;Gammaproteobacteria;Oceanospirillales;Halomonadaceae;Halomonas | 0.0031 |
| Patescibacteria;Microgenomatia;Candidatus Woesebacteria;uncultured Microgenomates group bacterium | 0.0029 |
| Verrucomicrobia;Verrucomicrobiae;Pedosphaerales;Pedosphaeraceae | 0.0028 |
| Bacteroidetes;Bacteroidia;Bacteroidales;Dysgonomonadaceae;Proteiniphilum | 0.0028 |
| Proteobacteria;Gammaproteobacteria;Betaproteobacteriales;Burkholderiaceae;Burkholderia-Caballeronia-Paraburkholderia | 0.0023 |
| Proteobacteria;Gammaproteobacteria;Methylococcales;Methylococcaceae;Methylococcus | 0.0022 |
| Proteobacteria;Alphaproteobacteria;Rhizobiales;Beijerinckiaceae;Methylocystis | 0.0019 |
| Firmicutes;Bacilli;Bacillales;Staphylococcaceae;Staphylococcus | 0.0018 |
| Firmicutes;Bacilli;Bacillales;Bacillaceae;Geobacillus | 0.0018 |
| Proteobacteria;Gammaproteobacteria;WD260 | 0.0015 |
| Bacteroidetes;Bacteroidia;Chitinophagales;Chitinophagaceae;Sediminibacterium | 0.0015 |
| Actinobacteria;Actinobacteria;Corynebacteriales;Corynebacteriaceae;Lawsonella | 0.0013 |
| Bacteroidetes;Bacteroidia;Sphingobacteriales;AKYH767;uncultured bacterium | 0.0012 |
| Bacteroidetes;Ignavibacteria;OPB56;uncultured Chlorobiales bacterium | 0.0012 |
| Actinobacteria;Actinobacteria;Corynebacteriales;Corynebacteriaceae;Corynebacterium 1 | 0.0011 |
| Proteobacteria;Alphaproteobacteria;Sphingomonadales;Sphingomonadaceae;Novosphingobium | 0.0011 |
| Actinobacteria;Actinobacteria;Streptomycetales;Streptomycetaceae;Streptomyces | 0.0010 |
| Firmicutes;Clostridia;Clostridiales;Peptostreptococcaceae;Romboutsia | 0.0010 |
| Proteobacteria;Gammaproteobacteria;Betaproteobacteriales;Burkholderiaceae;Achromobacter | 0.0009 |
| Proteobacteria;Gammaproteobacteria | 0.0008 |
| Proteobacteria;Deltaproteobacteria;Myxococcales;BIrii41;uncultured proteobacterium | 0.0007 |
| Proteobacteria;Gammaproteobacteria;Betaproteobacteriales;Burkholderiaceae;Comamonas | 0.0007 |
| Firmicutes;Bacilli;Bacillales;Alicyclobacillaceae;Alicyclobacillus | 0.0007 |
| Proteobacteria;Alphaproteobacteria;Sphingomonadales;Sphingomonadaceae;Sphingomonas | 0.0006 |
| Proteobacteria;Gammaproteobacteria;Xanthomonadales;Xanthomonadaceae;Luteimonas | 0.0006 |
| Actinobacteria;Actinobacteria;Micrococcales;Cellulomonadaceae;Cellulomonas | 0.0006 |
| Proteobacteria;Gammaproteobacteria;Pseudomonadales;Moraxellaceae;Moraxella | 0.0006 |
| Proteobacteria;Alphaproteobacteria;Rhodobacterales;Rhodobacteraceae;Paracoccus | 0.0006 |
| Proteobacteria;Gammaproteobacteria;Pseudomonadales;Moraxellaceae;Acinetobacter | 0.0005 |
| Acidobacteria;Subgroup 6;uncultured Anaeromyxobacter sp. | 0.0005 |
| Actinobacteria;Actinobacteria;Pseudonocardiales;Pseudonocardiaceae;Actinomycetospora | 0.0004 |
| Proteobacteria;Gammaproteobacteria;Enterobacteriales;Enterobacteriaceae;Pantoea | 0.0004 |
| Actinobacteria;Thermoleophilia;Solirubrobacterales;Solirubrobacteraceae;Conexibacter | 0.0004 |
| Proteobacteria;Alphaproteobacteria;Rhodobacterales;Rhodobacteraceae | 0.0004 |
| Actinobacteria;Actinobacteria;Frankiales;Geodermatophilaceae;Modestobacter | 0.0004 |
| Bacteroidetes;Ignavibacteria;Ignavibacteriales;Ignavibacteriaceae;Ignavibacterium | 0.0004 |
| Acidobacteria;Subgroup 6 | 0.0003 |
| Actinobacteria;Acidimicrobiia;IMCC26256; | 0.0002 |
| Proteobacteria;Gammaproteobacteria;Pseudomonadales;Pseudomonadaceae;Thiopseudomonas | 0.0002 |
| Proteobacteria;Alphaproteobacteria;Sphingomonadales;Sphingomonadaceae | 0.0002 |
| Bacteroidetes;Bacteroidia;Cytophagales;Spirosomaceae;Spirosoma | 0.0002 |
| Proteobacteria;Gammaproteobacteria;Betaproteobacteriales;Burkholderiaceae;Limnobacter | 0.0002 |
| Proteobacteria;Alphaproteobacteria;Rhizobiales;Beijerinckiaceae;1174-901-12 | 0.0001 |
