## Supplemental Table 2 for "Composition of the North American wood frog (*Rana sylvatica*) skin microbiome and seasonal variation in community structure"

**Supplementary Table 2** Bacterial taxa detected in field blanks and their mean relative abundance. Five field blanks were produced by exposing swabs to the open air and mimicking sampling, then following the outlined DNA extraction and sequencing procedures. Mean relative abundance values are given for genera where possible, or otherwise for most specific taxa known

| **Taxa** | **Mean Relative Abundance** |
| --- | --- |
| Actinobacteria;Actinobacteria;Micrococcales;Microbacteriaceae;Curtobacterium | 0.0839 |
| Proteobacteria;Gammaproteobacteria;Betaproteobacteriales;Burkholderiaceae;Burkholderia-Caballeronia-Paraburkholderia | 0.0708 |
| Proteobacteria;Gammaproteobacteria;Enterobacteriales;Enterobacteriaceae;Escherichia-Shigella | 0.0543 |
| Proteobacteria;Gammaproteobacteria;Pseudomonadales;Moraxellaceae;Acinetobacter | 0.0413 |
| Proteobacteria;Gammaproteobacteria;Betaproteobacteriales;Burkholderiaceae;Massilia | 0.0405 |
| Proteobacteria;Gammaproteobacteria;Pseudomonadales;Pseudomonadaceae;Pseudomonas | 0.0377 |
| Proteobacteria;Gammaproteobacteria;Betaproteobacteriales;Burkholderiaceae; | 0.0334 |
| Deinococcus-Thermus;Deinococci;Deinococcales;Deinococcaceae;Deinococcus | 0.0300 |
| Firmicutes;Bacilli;Bacillales;Alicyclobacillaceae;Alicyclobacillus | 0.0274 |
| Proteobacteria;Gammaproteobacteria;Pseudomonadales;Moraxellaceae;Alkanindiges | 0.0265 |
| Proteobacteria;Gammaproteobacteria;Immundisolibacterales;Immundisolibacteraceae;Immundisolibacter | 0.0255 |
| Tenericutes;Mollicutes;Acholeplasmatales;Acholeplasmataceae;Acholeplasma | 0.0232 |
| Proteobacteria;Gammaproteobacteria;Betaproteobacteriales;Burkholderiaceae;Duganella | 0.0230 |
| Proteobacteria;Gammaproteobacteria;Betaproteobacteriales;Burkholderiaceae;Ralstonia | 0.0211 |
| Proteobacteria;Alphaproteobacteria;Rhizobiales;Xanthobacteraceae; | 0.0206 |
| Proteobacteria;Alphaproteobacteria;Rhizobiales;Beijerinckiaceae;alphaI cluster | 0.0176 |
| Proteobacteria;Alphaproteobacteria;Sphingomonadales;Sphingomonadaceae;Sphingomonas | 0.0175 |
| Proteobacteria;Gammaproteobacteria;Xanthomonadales;Rhodanobacteraceae;Dyella | 0.0169 |
| Proteobacteria;Gammaproteobacteria;Betaproteobacteriales;Burkholderiaceae;Janthinobacterium | 0.0162 |
| Proteobacteria;Gammaproteobacteria;Xanthomonadales;Xanthomonadaceae;Stenotrophomonas | 0.0154 |
| Proteobacteria;Gammaproteobacteria;Betaproteobacteriales;Burkholderiaceae;Ramlibacter | 0.0131 |
| Bacteroidetes;Bacteroidia;Sphingobacteriales;Sphingobacteriaceae;Pedobacter | 0.0128 |
| Actinobacteria;Thermoleophilia;Gaiellales;uncultured; | 0.0109 |
| Proteobacteria;Alphaproteobacteria;Rhizobiales;Beijerinckiaceae;Methylocystis | 0.0105 |
| Proteobacteria;Alphaproteobacteria;Rhizobiales;Beijerinckiaceae;Methylobacterium | 0.0104 |
| Proteobacteria;Deltaproteobacteria;Desulfuromonadales;Geobacteraceae;Geothermobacter | 0.0092 |
| Firmicutes;Bacilli;Lactobacillales;Lactobacillaceae;Lactobacillus | 0.0092 |
| Bacteroidetes;Bacteroidia;Bacteroidales;Dysgonomonadaceae;Dysgonomonas | 0.0082 |
| Firmicutes;Bacilli;Lactobacillales;Streptococcaceae;Streptococcus | 0.0082 |
| Chloroflexi;P2-11E;uncultured bacterium;; | 0.0081 |
| Proteobacteria;Gammaproteobacteria;Pseudomonadales;Moraxellaceae;Psychrobacter | 0.0081 |
| Epsilonbacteraeota;Campylobacteria;Campylobacterales;Thiovulaceae;Sulfurimonas | 0.0080 |
| Atribacteria;JS1;uncultured bacterium;; | 0.0075 |
| Proteobacteria;Alphaproteobacteria;Rhodobacterales;Rhodobacteraceae;Paracoccus | 0.0075 |
| Proteobacteria;Alphaproteobacteria;Rhizobiales;Rhizobiaceae;Mesorhizobium | 0.0073 |
| Proteobacteria;Alphaproteobacteria;Rhizobiales;Beijerinckiaceae;1174-901-12 | 0.0068 |
| Bacteroidetes;Bacteroidia;Flavobacteriales;Weeksellaceae;Chryseobacterium | 0.0064 |
| Proteobacteria;Alphaproteobacteria;Sphingomonadales;Sphingomonadaceae;Polymorphobacter | 0.0062 |
| Actinobacteria;Actinobacteria;Frankiales;Sporichthyaceae;Candidatus Planktophila | 0.0062 |
| Actinobacteria;Actinobacteria;Corynebacteriales;Corynebacteriaceae;Lawsonella | 0.0054 |
| Proteobacteria;Gammaproteobacteria;Betaproteobacteriales;Chitinibacteraceae;Silvimonas | 0.0053 |
| Bacteroidetes;Bacteroidia;Flavobacteriales;Flavobacteriaceae;Flavobacterium | 0.0052 |
| Proteobacteria;Alphaproteobacteria;Rhodobacterales;Rhodobacteraceae; | 0.0050 |
| Acidobacteria;Acidobacteriia;Acidobacteriales;; | 0.0049 |
| Actinobacteria;Actinobacteria;Corynebacteriales;Corynebacteriaceae;Corynebacterium 1 | 0.0049 |
| Proteobacteria;Alphaproteobacteria;Sphingomonadales;Sphingomonadaceae;Rhizorhapis | 0.0046 |
| Armatimonadetes;Fimbriimonadia;Fimbriimonadales;Fimbriimonadaceae;uncultured bacterium | 0.0045 |
| Proteobacteria;Gammaproteobacteria;Betaproteobacteriales;Burkholderiaceae;Comamonas | 0.0044 |
| Actinobacteria;Actinobacteria;Micromonosporales;Micromonosporaceae; | 0.0044 |
| Actinobacteria;Actinobacteria;Micrococcales;Micrococcaceae;Rothia | 0.0043 |
| Bacteroidetes;Bacteroidia;Bacteroidales;Bacteroidaceae;Bacteroides | 0.0040 |
| Proteobacteria;Deltaproteobacteria;Myxococcales;Sandaracinaceae;uncultured | 0.0039 |
| Firmicutes;Clostridia;Clostridiales;Ruminococcaceae;Ethanoligenens | 0.0037 |
| Tenericutes;Mollicutes;EUB33-2;uncultured bacterium; | 0.0036 |
| Proteobacteria;Gammaproteobacteria;Enterobacteriales;Enterobacteriaceae; | 0.0036 |
| Firmicutes;Bacilli;Bacillales;Bacillaceae;Bacillus | 0.0035 |
| Actinobacteria;Acidimicrobiia;IMCC26256;; | 0.0034 |
| Actinobacteria;Actinobacteria;Micrococcales;Micrococcaceae;Kocuria | 0.0033 |
| Planctomycetes;Planctomycetacia;Pirellulales;Pirellulaceae;Pirellula | 0.0032 |
| Proteobacteria;Alphaproteobacteria;Caulobacterales;Caulobacteraceae;Brevundimonas | 0.0031 |
| Acidobacteria;Acidobacteriia;Acidobacteriales;uncultured; | 0.0030 |
| Actinobacteria;Actinobacteria;Micrococcales;Micrococcaceae;Micrococcus | 0.0030 |
| Bacteroidetes;Bacteroidia;Chitinophagales;Chitinophagaceae;uncultured | 0.0030 |
| Proteobacteria;Alphaproteobacteria;;; | 0.0030 |
| Bacteroidetes;Bacteroidia;Chitinophagales;Chitinophagaceae;Cnuella | 0.0030 |
| Actinobacteria;Actinobacteria;Corynebacteriales;Mycobacteriaceae;Mycobacterium | 0.0029 |
| Verrucomicrobia;Verrucomicrobiae;Chthoniobacterales;Chthoniobacteraceae;Chthoniobacter | 0.0029 |
| Firmicutes;Clostridia;Clostridiales;Family XI;Finegoldia | 0.0028 |
| Proteobacteria;Alphaproteobacteria;Rhodospirillales;uncultured;uncultured Rhodospirillaceae | 0.0028 |
| Actinobacteria;Actinobacteria;Micrococcales;Intrasporangiaceae; | 0.0027 |
| Actinobacteria;Actinobacteria;Micrococcales;Microbacteriaceae;Agromyces | 0.0027 |
| Proteobacteria;Alphaproteobacteria;Acetobacterales;Acetobacteraceae;Acidiphilium | 0.0026 |
| Chlamydiae;Chlamydiae;Chlamydiales;Parachlamydiaceae;uncultured | 0.0024 |
| Actinobacteria;Actinobacteria;Propionibacteriales;Nocardioidaceae;Marmoricola | 0.0023 |
| Firmicutes;Bacilli;Bacillales;Staphylococcaceae;Staphylococcus | 0.0022 |
| Proteobacteria;Alphaproteobacteria;Sphingomonadales;Sphingomonadaceae;uncultured | 0.0021 |
| Acidobacteria;Acidobacteriia;Acidobacteriales;uncultured Acidobacteriaceae bacterium; | 0.0021 |
| Proteobacteria;Gammaproteobacteria;Xanthomonadales;Xanthomonadaceae;Arenimonas | 0.0020 |
| Proteobacteria;Alphaproteobacteria;Sphingomonadales;Sphingomonadaceae; | 0.0018 |
| Verrucomicrobia;Verrucomicrobiae;Opitutales;Opitutaceae;Opitutus | 0.0016 |
| Bacteroidetes;Bacteroidia;Cytophagales;Hymenobacteraceae;Hymenobacter | 0.0016 |
| Proteobacteria;Gammaproteobacteria;Betaproteobacteriales;Hydrogenophilaceae;Thiobacillus | 0.0015 |
| Verrucomicrobia;Verrucomicrobiae;Verrucomicrobiales;Rubritaleaceae;Luteolibacter | 0.0015 |
| Actinobacteria;Actinobacteria;Micrococcales;Microbacteriaceae; | 0.0015 |
| Proteobacteria;Gammaproteobacteria;Enterobacteriales;Enterobacteriaceae;Pantoea | 0.0015 |
| Proteobacteria;Alphaproteobacteria;Sphingomonadales;Sphingomonadaceae;Altererythrobacter | 0.0015 |
| Proteobacteria;Gammaproteobacteria;Betaproteobacteriales;Methylophilaceae;Methylotenera | 0.0015 |
| Actinobacteria;Actinobacteria;Micrococcales;; | 0.0014 |
| Proteobacteria;Gammaproteobacteria;Betaproteobacteriales;Burkholderiaceae;Variovorax | 0.0013 |
| Bacteroidetes;Bacteroidia;Sphingobacteriales;Sphingobacteriaceae;Mucilaginibacter | 0.0012 |
| Actinobacteria;Actinobacteria;Corynebacteriales;Nocardiaceae;Rhodococcus | 0.0012 |
| Proteobacteria;Gammaproteobacteria;Betaproteobacteriales;Burkholderiaceae;Rhodoferax | 0.0012 |
| Proteobacteria;Gammaproteobacteria;Gammaproteobacteria Incertae Sedis;Unknown Family;Acidibacter | 0.0011 |
| Chloroflexi;Anaerolineae;Caldilineales;Caldilineaceae;uncultured | 0.0011 |
| Proteobacteria;Gammaproteobacteria;Xanthomonadales;Rhodanobacteraceae;Rhodanobacter | 0.0011 |
| Actinobacteria;Actinobacteria;Propionibacteriales;Nocardioidaceae;Nocardioides | 0.0010 |
| Actinobacteria;Thermoleophilia;Solirubrobacterales;67-14;metagenome | 0.0010 |
| Proteobacteria;Gammaproteobacteria;Pseudomonadales;Pseudomonadaceae;Thiopseudomonas | 0.0009 |
| Firmicutes;Bacilli;Bacillales;Family XI;Gemella | 0.0009 |
| Actinobacteria;Actinobacteria;Micrococcales;Dermabacteraceae;Brachybacterium | 0.0009 |
| Acidobacteria;Holophagae;Subgroup 7;; | 0.0009 |
| Proteobacteria;Gammaproteobacteria;Betaproteobacteriales;Burkholderiaceae;Schlegelella | 0.0009 |
| Proteobacteria;Deltaproteobacteria;Desulfarculales;Desulfarculaceae;uncultured | 0.0009 |
| Proteobacteria;Alphaproteobacteria;Rhizobiales;Xanthobacteraceae;uncultured | 0.0009 |
| Proteobacteria;Alphaproteobacteria;Sphingomonadales;Sphingomonadaceae;Sphingobium | 0.0009 |
| Bacteroidetes;Bacteroidia;Bacteroidales;Prevotellaceae;Prevotella 7 | 0.0008 |
| Actinobacteria;Actinobacteria;Micrococcales;Dermacoccaceae;Dermacoccus | 0.0008 |
| Actinobacteria;Thermoleophilia;Solirubrobacterales;67-14;uncultured Solirubrobacter sp. | 0.0008 |
| Bacteroidetes;Bacteroidia;Cytophagales;Spirosomaceae;Dyadobacter | 0.0008 |
| Actinobacteria;Actinobacteria;Corynebacteriales;Tsukamurellaceae;Tsukamurella | 0.0008 |
| Actinobacteria;Actinobacteria;Pseudonocardiales;Pseudonocardiaceae;Actinomycetospora | 0.0008 |
| Proteobacteria;Alphaproteobacteria;Caulobacterales;Caulobacteraceae;Caulobacter | 0.0007 |
| Proteobacteria;Alphaproteobacteria;Rhizobiales;Beijerinckiaceae;Roseiarcus | 0.0007 |
| Proteobacteria;Alphaproteobacteria;Rhizobiales;Xanthobacteraceae;Bradyrhizobium | 0.0007 |
| FBP;uncultured bacterium;;; | 0.0006 |
| Proteobacteria;Gammaproteobacteria;Pseudomonadales;Moraxellaceae;Enhydrobacter | 0.0006 |
| Proteobacteria;Deltaproteobacteria;Myxococcales;Polyangiaceae;Pajaroellobacter | 0.0006 |
| Proteobacteria;Gammaproteobacteria;Immundisolibacterales;Immundisolibacteraceae; | 0.0006 |
| Actinobacteria;Actinobacteria;Frankiales;Geodermatophilaceae;Blastococcus | 0.0005 |
| Proteobacteria;Alphaproteobacteria;Rickettsiales;SM2D12;unidentified marine bacterioplankton | 0.0005 |
| Actinobacteria;Actinobacteria;Micrococcales;Brevibacteriaceae;Sediminivirga | 0.0005 |
| Proteobacteria;Gammaproteobacteria;Xanthomonadales;Xanthomonadaceae;Pseudoxanthomonas | 0.0005 |
| Proteobacteria;Gammaproteobacteria;Betaproteobacteriales;Nitrosomonadaceae;MND1 | 0.0005 |
| Actinobacteria;Actinobacteria;Micromonosporales;Micromonosporaceae;Luedemannella | 0.0005 |
| Proteobacteria;Alphaproteobacteria;Rhizobiales;Beijerinckiaceae;Microvirga | 0.0005 |
| Proteobacteria;Gammaproteobacteria;Betaproteobacteriales;A21b; | 0.0005 |
| Proteobacteria;Gammaproteobacteria;Betaproteobacteriales;Burkholderiaceae;Achromobacter | 0.0005 |
| Proteobacteria;Gammaproteobacteria;Betaproteobacteriales;Burkholderiaceae;Hydrogenophaga | 0.0004 |
| Proteobacteria;Alphaproteobacteria;Rhizobiales;Rhizobiaceae;Allorhizobium-Neorhizobium-Pararhizobium-Rhizobium | 0.0004 |
| Actinobacteria;Actinobacteria;Micromonosporales;Micromonosporaceae;Micromonospora | 0.0004 |
| Proteobacteria;Deltaproteobacteria;Bdellovibrionales;Bacteriovoracaceae;Peredibacter | 0.0004 |
| Bacteroidetes;Bacteroidia;Chitinophagales;Chitinophagaceae;Ferruginibacter | 0.0004 |
| Actinobacteria;Actinobacteria;Micrococcales;Dermacoccaceae;Kytococcus | 0.0004 |
| Actinobacteria;Actinobacteria;Frankiales;; | 0.0004 |
| Actinobacteria;Actinobacteria;Micrococcales;Dermatophilaceae;Mobilicoccus | 0.0004 |
| Actinobacteria;Actinobacteria;Micrococcales;Brevibacteriaceae;Brevibacterium | 0.0004 |
| Firmicutes;Bacilli;Bacillales;Staphylococcaceae;Salinicoccus | 0.0003 |
| Proteobacteria;Alphaproteobacteria;Rhodobacterales;Rhodobacteraceae;Rhodobacter | 0.0003 |
| Actinobacteria;Thermoleophilia;Solirubrobacterales;Solirubrobacteraceae;Conexibacter | 0.0003 |
| Actinobacteria;Actinobacteria;Streptomycetales;Streptomycetaceae;Streptomyces | 0.0003 |
| Proteobacteria;Gammaproteobacteria;Aeromonadales;Aeromonadaceae;Aeromonas | 0.0003 |
| Proteobacteria;Gammaproteobacteria;Alteromonadales;Alteromonadaceae;Rheinheimera | 0.0003 |
| Proteobacteria;Deltaproteobacteria;Bdellovibrionales;Bdellovibrionaceae;Bdellovibrio | 0.0003 |
| Actinobacteria;Actinobacteria;Kineosporiales;Kineosporiaceae; | 0.0003 |
| Proteobacteria;Gammaproteobacteria;Betaproteobacteriales;Neisseriaceae;Neisseria | 0.0003 |
| Proteobacteria;Alphaproteobacteria;Rhizobiales;Methyloligellaceae;uncultured | 0.0003 |
| Proteobacteria;Gammaproteobacteria;Alteromonadales;Shewanellaceae;Shewanella | 0.0003 |
| Proteobacteria;Gammaproteobacteria;Betaproteobacteriales;SC-I-84;metagenome | 0.0003 |
| Proteobacteria;;;; | 0.0003 |
| Firmicutes;Clostridia;Clostridiales;Family XI;Anaerococcus | 0.0002 |
| Acidobacteria;Acidobacteriia;Acidobacteriales;Acidobacteriaceae (Subgroup 1);uncultured | 0.0002 |
| Actinobacteria;Actinobacteria;Frankiales;Geodermatophilaceae;Modestobacter | 0.0002 |
| Deinococcus-Thermus;Deinococci;Deinococcales;Trueperaceae;Truepera | 0.0002 |
| Acidobacteria;Subgroup 6;;; | 0.0002 |
| Bacteroidetes;Bacteroidia;Cytophagales;Spirosomaceae;Emticicia | 0.0002 |
| Actinobacteria;Thermoleophilia;Gaiellales;Gaiellaceae;Gaiella | 0.0002 |
| Planctomycetes;Planctomycetacia;Gemmatales;Gemmataceae;Gemmata | 0.0002 |
| Proteobacteria;Gammaproteobacteria;Xanthomonadales;Xanthomonadaceae;Lysobacter | 0.0002 |
| Proteobacteria;Alphaproteobacteria;Rhizobiales;Beijerinckiaceae;Bosea | 0.0001 |
| Proteobacteria;Alphaproteobacteria;Rhizobiales;Beijerinckiaceae; | 0.0001 |
| Proteobacteria;Alphaproteobacteria;Dongiales;Dongiaceae;Dongia | 0.0001 |
| Proteobacteria;Gammaproteobacteria;Betaproteobacteriales;SC-I-84;uncultured Nitrosomonadaceae | 0.0001 |
| Bacteroidetes;Bacteroidia;Chitinophagales;Chitinophagaceae;Niastella | 0.0001 |
| Proteobacteria;Gammaproteobacteria;Steroidobacterales;Steroidobacteraceae;Steroidobacter | 0.0000 |
